# Restoring functional D2- to D1-neuron correspondence enables goal-directed action control in long-lived striatal circuits

**DOI:** 10.1101/2022.09.14.508004

**Authors:** Jesus Bertran-Gonzalez, Caroline Dinale, Miriam Matamales

## Abstract

Multidisciplinary evidence suggests that instrumental performance is governed by two major forms of behavioural control: goal-directed and autonomous processes. Brain-state abnormalities affecting the striatum, such as ageing, often shift control towards autonomous—habit-like—behaviour, although the neural mechanisms responsible for this shift remain unknown. Here, combining instrumental conditioning with cell-specific functional mapping and manipulation in striatal neurons, we explored strategies that invigorate goal-directed action capacity in aged mice. In animals performing instrumental actions, D2- and D1-neurons of the aged striatum were engaged in a characteristically counterbalanced manner, something that related to the propensity to express autonomous behaviour. Long-lasting, cell-specific desensitisation of D2-neurons in aged transgenic mice recapitulated the uneven D2-to D1-neuron functional correspondence observed in young mice, an effect that enabled successful goal-directed action. Our findings contribute to the understanding of the neural bases of behavioural control and propose neural system interventions that enhance cognitive functioning in habit-prone brains.

## Introduction

Two alternative behavioural control processes have been identified as fundamental drivers of instrumental action (Dickinson, 1985). The first controller—known as goal-directed, cognitive, teleological or model-based, amongst other names—strongly relies on the association between actions (A) and the outcomes (O) they produce (A→O contingency). As such, actions are truly purposeful, for they are sensitive to both the causal relationship to their consequences and the value of those consequences (Balleine and Dickinson, 1998). The second controller—known as habit, autonomous or model-free—is thought to depend on strong associative links formed between antecedent stimuli and the responses that lead to outcomes (S→R association), prompting actions that are performed independently of their consequences (Dickinson, 1994). Several dual-process theories propose a variety of ways by which one and/or the other controller assumes governance of behaviour, including transitional (Dickin-son, 1985, 1994), arbitrary (Daw et al., 2005) and collaborative accounts (Balleine and Dezfouli, 2019; Dezfouli and Balleine, 2013; Dickinson, 2012).

A first group of factors influencing whether goal-directed or autonomous controllers take over behaviour relates to the *conditions of learning*. For example, goal-direction can be protected from autonomy if the A→O contingency is readily accessible (Dickinson, 1985; Perez and Dickinson, 2020), or if the complexity of the task increases, such as when multiple A→O contingencies are learned simultaneously (Colwill and Rescorla, 1985; Kosaki and Dickinson, 2010). Thus, while it is often assumed that behaviour, once it is rendered autonomous, becomes inflexible and insensitive to change, even extensively-trained habits have been rendered goal-directed by manipulations of the conditions of learning (Trask et al., 2020). On the other hand, a second group of factors determining predominance of one or the other process relates to the *state of the brain* at the moment of learning. Both temporary processes such as sensitisation to drugs (Nelson and Killcross, 2006), working memory loads and stress (Otto et al., 2013), as well as longer-lived states such as drug addiction (Ostlund and Balleine, 2008), stroke (Cognat et al., 2010; Habib, 2004) and ageing (Eppinger et al., 2013; Ito et al., 2021; Mata et al., 2010; de Wit et al., 2014), have been reported to bias goal-directed behavioural command towards an—often detrimental—autonomous control. These conditions share the property of interfering either directly or indirectly with the function of the dorsal striatum, in particular with its medial portion (i.e., the dorsomedial striatum; DMS), an area that is centrally involved in encoding the A→O associative processes supporting goal-directed action (Balleine, 2019).

Within the DMS, research in the last few years has started to clarify the role played by its two main neuronal components—the D1- and D2-expressing spiny projection neurons; D1- and D2-SPNs—in the integration of goal-directed learning (Balleine et al., 2021). Evidence mounts for widespread reciprocal modulations amongst SPNs operating across the DMS and beyond (Burke et al., 2017; Ponzi and Wickens, 2013), and with those, more sophisticated appreciations of the actual functional histoarchitectural logic of the striatum emerge (Tinterri et al., 2018). Recent findings in our lab highlight the importance of the regulatory transmodulations established locally between D2- and D1-SPNs for the re-writing of previously established goal-directed memories, a necessary process for producing behavioural change (Matamales et al., 2020). Our views identify a functional binary mosaic in the striatum by which the proportion of engaged D2- and D1-SPNs along with their spatial confluence are the key phenomena predicting the learning capacities of a particular striatal domain (Matamales et al., 2020).

In the case of ageing, brains are proposed to undergo deep compensatory neural adaptations to various fronts of structural and neuronal decline (Park and Reuter-Lorenz, 2009), which result in characteristic overactivations of frontal cortical and fronto-medial corticostriatal circuitries (Matamales et al., 2017; Park and Reuter-Lorenz, 2009; Zhang et al., 2017). Such adaptations encompass circuitry around the DMS (Matamales et al., 2016, 2017) and so likely relate to some, if not all, of the deficits in A→O encoding commonly observed in ageing (Matamales et al., 2016; O’Callaghan et al., 2019; de Wit et al., 2014). However, the way ageing-related circuit reorganisations affect the functional distributions of D1- and D2-SPNs in the striatum has never been studied. Collectively, whether ageing brings about a state by which adaptations in the local striatal circuitry favour autonomous control, and whether such state can be internally altered to promote goal-direction, remain open questions.

Here, we sought to determine the origin of autonomous dominance in ageing by disambiguating the influence of factors relating to the *conditions of learning* from those relating to the *state of the brain*. We found that while diverse manipulations of the conditions of learning failed to enable goal-directed control in aged mice, the aged striatum presented characteristic functional features involving counterbalanced proportions of D2- and D1-SPNs in the DMS that suggested a state of plasticity stillness. We showed that recreating the functional cellular environment observed in the DMS of young mice through long-lasting, circuit-specific adaptations in aged transgenic mice proved successful at enabling their ability to integrate A→O contingencies, something that promoted vigorous goal-directed action.

## Results

### Instrumental performance in aged mice may not be fit for appropriate A→O encoding

To investigate whether young and aged mice presented similar opportunity to integrate A→O contingencies on training, we studied details of their instrumental performance during simple action (A; lever press) → outcome (O; grain food pellet) arrangements. We trained mildly food restricted mice on an increasing random ratio (RR) schedule of reinforcement, where the number of presses required to obtain outcomes increased as training progressed (from constant reinforcement, CRF, to random ratio 10, RR10; Fig. 1a). We monitored the formation of action sequences by parsing behavioural data into initiation, execution and termination categories, as done previously (Matamales et al., 2017) (Fig. 1a). While both young and aged groups significantly escalated their performance as the A→O ratio requirement increased (Fig. 1b), aged mice showed significantly reduced rates towards the end of training (Supplementary Table 1). We examined the spontaneous structuring of behaviour by comparing performance between days 7 and 15 of training. We found that young mice significantly increased the number of execution elements per sequence, whereas aged mice kept a similar number throughout training (Fig. 1c, Supplementary Table 1). Consistently, while the sequence duration in young mice extended across training, it remained unchanged in aged mice (Fig. 1d, Supplementary Table 1). Event-time diagrams provided a visual representation of this effect (Fig. 1e and Extended Data Fig. 1): from days 7 to 15, young mice structured their behaviour into defined action sequences with clearly identifiable initiation, execution and termination elements, whereas aged mice showed no signs of consistent sequence structuring (Fig. 1e and Extended Data Fig. 1).

**Fig. 1.**
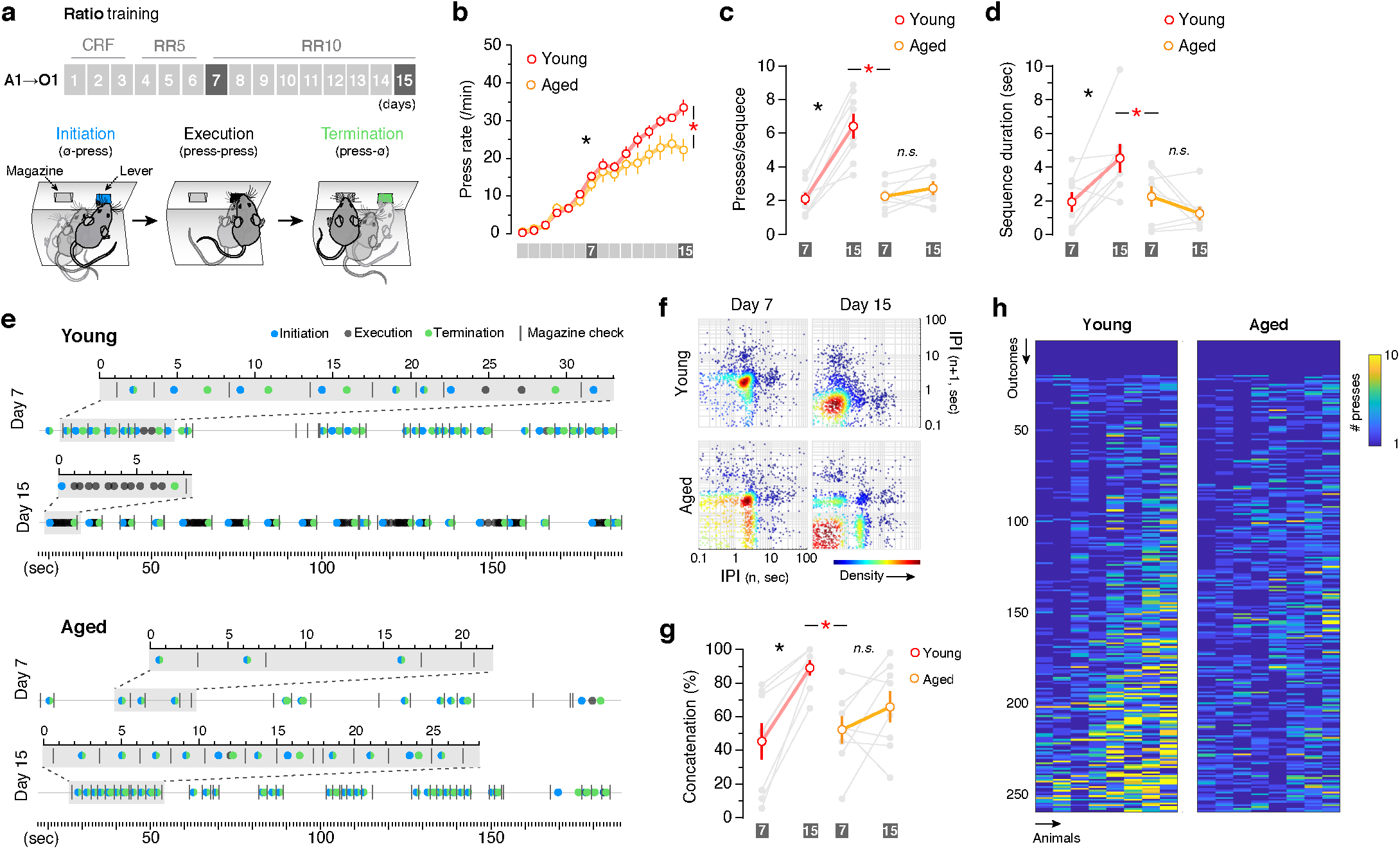
Aged mice show impaired sequential action when trained on single action → outcome relationships. **a**, Top: mice were trained on a single action (A1) → outcome (O1) contingency under continuous reinforcement (CRF) followed by increasing random ratio (RR) reinforcement schedules. Bottom: performance was distributed over initiation, execution and termination categories based on lever press and magazine entry interspersing. **b**, Lever presses per minute (press rate) displayed throughout instrumental training. **c**-**d**, Average presses per sequence and sequence duration (e) by young and aged mice on days 7 and 15. **e**, Event-time diagrams showing linearly organised lever press sequences produced by one young and one aged mouse on pre- (day 7) and post-sequence (day 15) stages of training. Shaded segments (grey) are amplified sequences (seconds). See Extended Data Fig. 1 for diagrams in all animals. **f**, Return maps showing the temporal relationships between consecutive lever presses at different stages of A→O training. Performance from all animals of each group at the indicated day is shown. Each dot corresponds to a pair of consecutive IPIs and includes the time delay of each press to its preceding (n [x]) and succeeding (n + 1 [y]) press in the sequence (in seconds [log_10_]). Pseudocolouring of spatial densities highlights data clustering. **g**, Percentage of lever press concatenation defined as number of sequences with ≥ 2 presses over total. **h**, Heatmap showing the concatenation extent of lever press sequences preceding each outcome, organised chronologically from start to end of training. Grey traces in d, e and g are individual mice. Data are mean ± SEM; 8 mice per group; *, overall/simple effect (black) and interaction (red). *n*.*s*., not significant (Supplementary Table 1).

We further explored action structure deficits by mapping the temporal relationships between consecutive lever presses (inter press intervals, IPIs) at different phases of training (Fig. 1f). This provided a visual readout of the behavioural patterns that emerged in each group during training (Jin et al., 2014): lever press performance in young mice accelerated smoothly from a defined cluster on day 7 (∼2 sec IPIs) to a stable temporal space (∼0.4 sec IPIs) on day 15 (Fig. 1f, top). Aged mice, however, transitioned from a similar performance on day 7 (∼2 sec) to a scattered pattern of performance on day 15, with a main large cluster (red) involving fast and very fast responses (0.1-0.5 sec IPIs) and a second cluster (green) involving responses that were equal to those in day 7 (∼2 sec) (Fig. 2f, bottom). Such pattern mostly emerged from non-concatenated action chains, as revealed by the limited increase in the proportion of sequences bearing 2 or more presses in aged as opposed to young mice (Fig. 1g and h, Supplementary Table 1). These results show a lack of sequential structure in aged mice that could interfere with the appropriate behaviour-reward correlation proposed to support the development of goal-direction (Dickinson, 1985). Therefore, in these conditions, young and aged mice displayed dissimilar opportunities to integrate the A→O contingency necessary for goal-directed control.

**Fig. 2.**
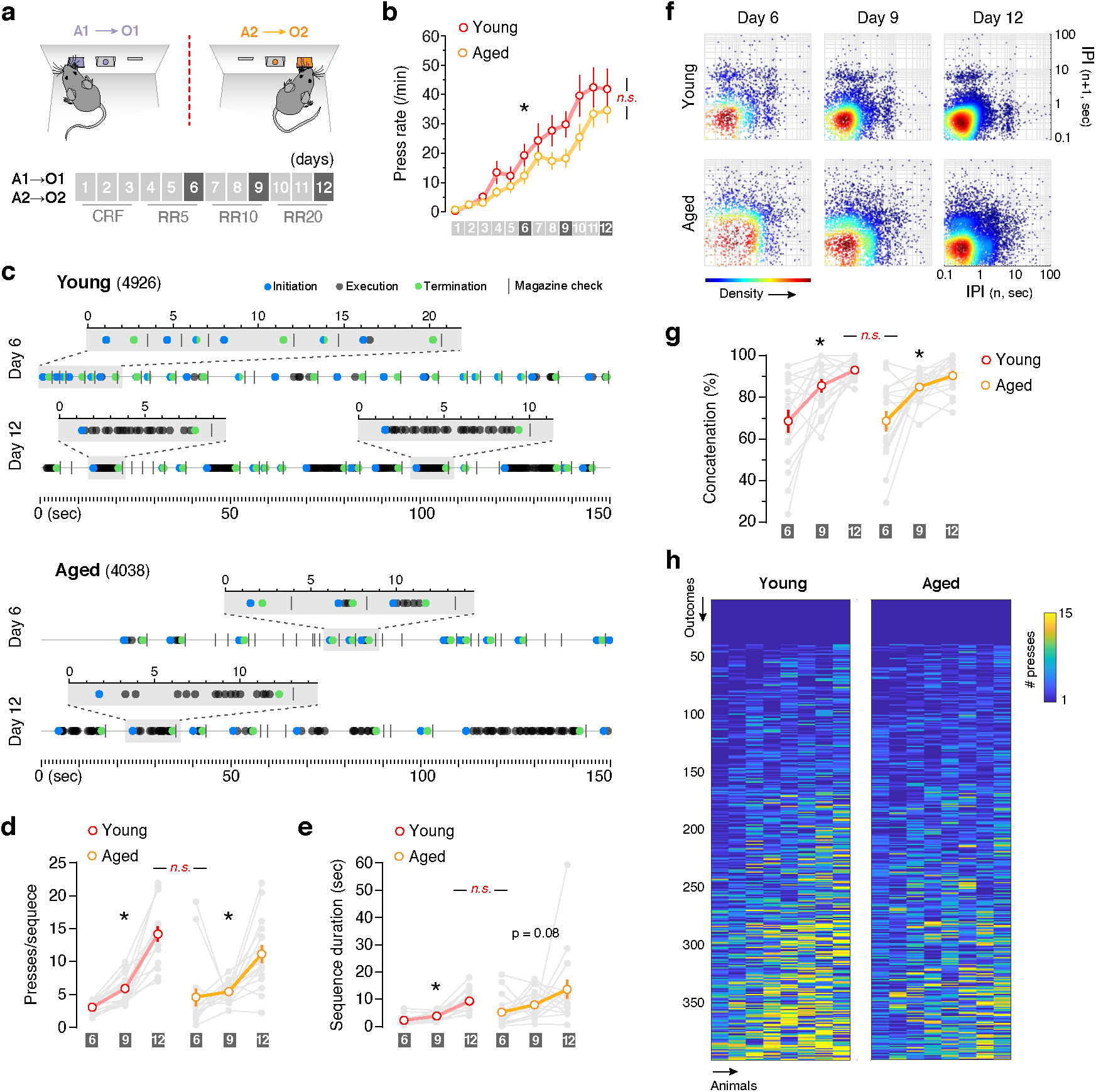
Sequential action is preserved in aged mice when two A→O contingencies are trained side by side. **a**, Mice were trained on two different action → outcome contingencies (A1→O1; A2→O2) under continuous reinforcement (CRF) followed by increasing random ratio (RR) reinforcement schedules. Contingencies were trained in separate sessions each day (red dashed line). **b**, Lever presses per minute (press rate) displayed throughout instrumental training (averaged A1 and A2; 8 mice per group). **c**, Event-time diagrams showing linearly organised lever press sequences produced in one lever by one young and one aged mouse on pre- (day 6) and post-sequence (day 12) stages of training. Shaded segments (grey) are amplified sequences (seconds). See Extended Data Fig. 1 for diagrams in all animals. **d**-**e**, Average presses per sequence (d) and sequence duration by young and aged mice on days 6 and 12. **f**, Return maps showing the temporal relationships between consecutive lever presses at different stages of A→O training. Performance from all animals of each group at the indicated day is shown. Each dot corresponds to a pair of consecutive IPIs and includes the time delay of each press to its preceding (n [x]) and succeeding (n + 1 [y]) press in the sequence (in seconds [log_10_]). Pseudocolouring of spatial densities highlights data clustering. **g**, Percentage of lever press concatenation defined as number of sequences with ≥ 2 presses over total. **h**, Heatmap showing the concatenation extent of lever press sequences preceding each outcome, organised chronologically from start to end of training. Data in d-h are pooled A1→O1 and A2→O2 performance. Data are mean ± SEM. Grey traces in d, e and g are individual mice (8 mice, 2 sessions per mouse). *, overall/simple effect (black) and interaction (red). *n*.*s*., not significant (Supplementary Table 1).

### Training on multiple A→O contingencies promotes adequate action structure in aged mice

We next sought strategies that improve the conditions of learning in aged mice in an attempt to equate their goal-directed control capacity to that projected in young mice. We explored the effects of adding a second A→O contingency on performance during escalating RR schedules, something that has been shown effective at preserving goal-direction throughout extended training on interval schedules in rats (Kosaki and Dickinson, 2010; Trask et al., 2020). We trained animals to two distinct A→O contingencies, A1→O1 and A2→O2, each involving a different lever (right or left) and a different outcome (grain or purified food pellets, see Supplementary Methods) (Fig. 2a). We found a trend for reduced overall lever press performance in aged mice that failed to reach statistical significance (Fig. 2b, Supplementary Table 1). Analysis of behavioural structure showed that, in contrast to single A→O training, aged mice trained on two A→O contingencies were able to develop action sequences throughout training, with clearly defined sequence elements (Fig. 2c, Extended Data Fig. 2) and similar rate and duration to those produced by young controls (Fig. 2d and e, Supplementary Table 1). Analysis of IPI distribution revealed equivalent behavioural patterns across age groups, with both young and aged mice showing similar clustering of IPIs at ∼0.3 sec by day 12 (Fig. 2f). These matching patterns were supported by similar concatenation rates in both groups—aged mice now showed a clear ability to produce behavioural chains throughout training (Fig. 2g and h, Supplementary Table 1).

Based on the known differences in performance resulting from ratio and interval schedules (Dickinson et al., 1983), we sought to explore the effects of training two A→O contingencies on sequence performance under increasing random interval (RI) schedules of reinforcement (Extended Data Fig. 3a). Similar to the performance observed in aged mice during single A→ O training (cf. Fig 1), this training induced virtually no sequence structure in either age group (Extended Data Fig. 3b-d, Extended Data Fig. 4). This suggested that, in contrast to findings in rats (Kosaki and Dickinson, 2010; Trask et al., 2020), RI schedules in mice may not immediately promote correlated A→O encoding that is competent for goal-directed control, even when multiple A→O contingencies are trained side by side. On RR schedules, however, our results showed that addition of a second A→O contingency largely improved sequence structure in aged mice; both groups now showed a level of performance in training that appeared propitious for the development of goal-directed control.

### Aged mice show resilient autonomous control

Our data suggested that training a second A→O contingency benefits action sequence performance in aged mice moderately trained to ratio schedules, an effect that could facilitate an adequate encoding of A→O contingencies and, with it, the engagement of goal-directed control in ageing (Daw et al., 2005; Dickinson, 1985; Eppinger et al., 2013; Kosaki and Dickinson, 2010; Trask et al., 2020). We sought to address this specifically by exposing the mice pretrained on multiple A→O contingencies to an outcome-specific devaluation test, which evaluated the effect of satiety to one or the other outcome on the choice between the two trained instrumental actions (Balleine and Dickinson, 1998) (Fig. 3a). If the mice correctly encoded these contingencies (i.e., A1→O1 and A2→O2), satiety on one of the outcomes (e.g., O1) should reduce performance of the action associated with this particular outcome (A1; devalued) relative to performance of the other, still valued action (A2; valued). To test this, mice were given unrestricted access to either O1 or O2 for 1h followed by a choice extinction test, in which—for the first time—both levers were available but no outcome was delivered (Fig. 3b). We found that while young mice correctly adjusted their performance towards the valued action, aged mice showed no clear preference on choice (Fig. 3c, Supplementary Table 1), indicating that, despite displaying appropriate action sequence performance during training (c.f. Fig. 2), aged mice still showed deficits in goal-directed encoding. We next investigated whether additional signs of goal-direction were observable through detailed analysis of lever press performance during the extinction choice tests. We found that young mice vigorously engaged with the valued-paired lever intermittently, usually after periods of explorative performance on the devalued-paired lever, whereas aged mice showed a homogeneous distribution of performance on both levers throughout the session, with no signs of vigorous alternation (Fig. 3d and e). In parallel, using a new cohort of animals, we explored the possibility that training aged mice on both A→O contingencies interspersedly during the same session (instead of sequentially in separate sessions as done thus far) could encourage better A→O encoding, a requirement deemed important to facilitate goal-directed control in rats (Kosaki and Dickinson, 2010; Trask et al., 2020). Aged mice trained in this way readily acquired lever press performance (Extended Data Fig. 5a, Supplementary Table 1), although they continued to show no preference towards valued- or devalued-paired levers on test (Fig. 3f, Supplementary Table 1). Again, interspersed training of the two A→O contingencies did not improve vigorous responding on any of the levers during the extinction choice tests in aged mice (Fig. 3g, Extended Data Fig. 5b), nor did training the same contingencies on interval schedules, which showed no signs of goal-direction in any of the age groups (Extended Data Fig. 6, Supplementary Table 1).

**Fig. 3.**
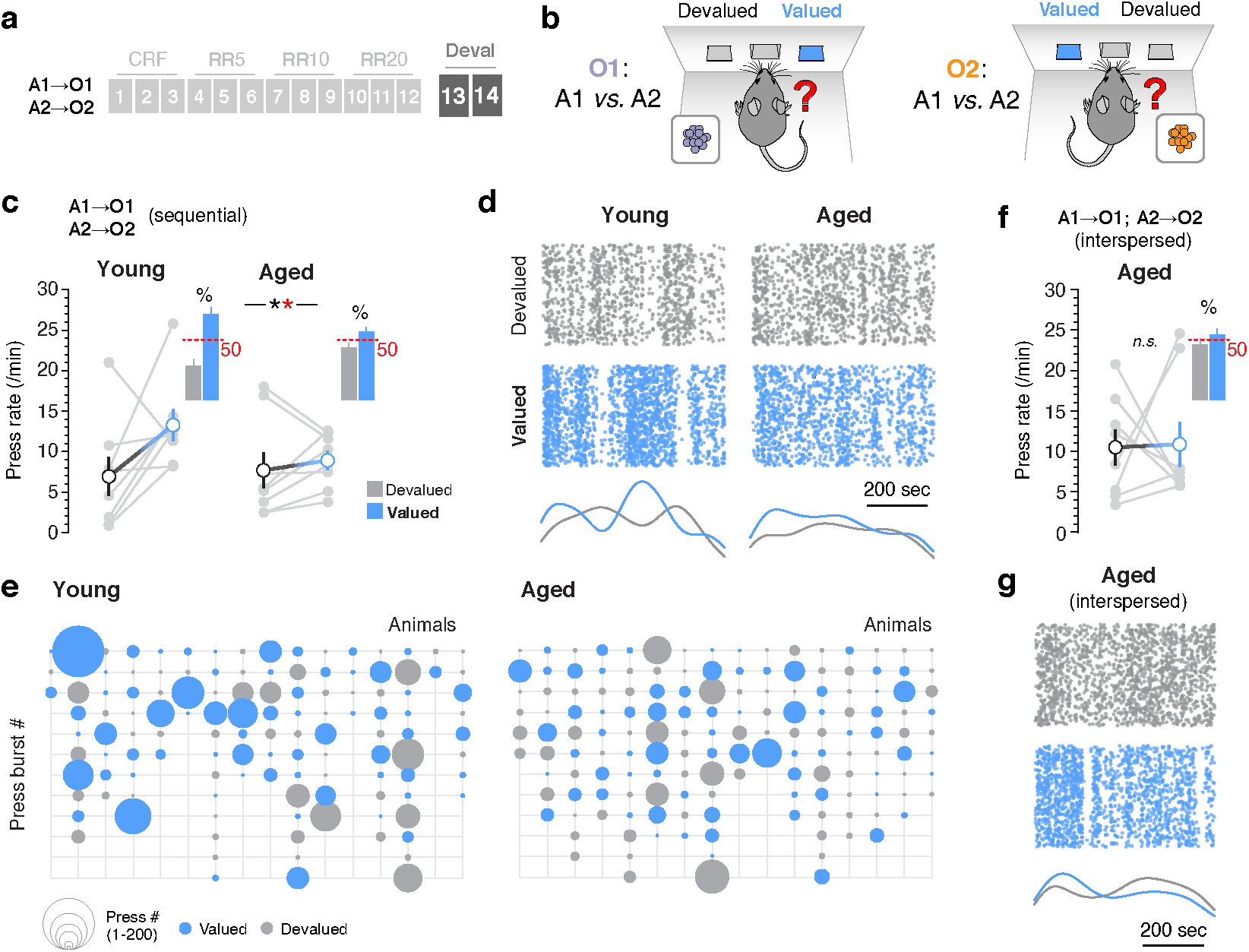
Goal-direction holds off in aged mice. **a**, Two separate action → outcome contingencies (A1→O1 and A2→O2) were trained either sequentially (in separate sessions; panels c-e) or interspersedly (within the same sessions; panels f-g) at continuous (CRF) and increasing random ratio (RR5-RR20) reinforcement schedules. **b**, assessment of A→O contingency learning through outcome-specific devaluation tests (Deval), where mice were sated for 1 hr on one or the other outcome (i.e., O1 or O2) prior to a choice test (A1 versus A2). **c** and **f**, Press rates on the lever coding for the sated (devalued, grey) and nonsated (valued, blue) outcomes during the choice tests after sequential (c) and interspersed (f) A→O training. Pairs of data points (light grey) are individual trials (each mouse was exposed to one trial per day on two consecutive days; see Supplementary Methods for details). Insets show the percentage of presses devoted to valued-paired or devalued-paired levers over the total presses in the tests (dashed red line denotes 50%). **d** and **g**, Scatter plot of all lever presses performed by the full cohort of young and aged mice during the 10-min choice tests after sequential (d) and interspersed (g) A→O training. Performance is distributed over devalued- (grey) and valued- (blue) paired levers. Corresponding Kernel density curves of each lever type are shown at the bottom. **e**, Bubble plot representing the response vigour exerted on devalued- (grey) and valued- (blue) paired levers throughout the choice test. Each column represents performance by one mouse in one session (two columns per mouse). Press bursts are defined as all performance within a 20 second interval, and are organised chronologically from top to bottom. Data are mean ± SEM or mean + SEM; 8 mice per group; *, overall/simple effect (black) and interaction (red). *n*.*s*., not significant (Supplementary Table 1).

### Aged SPN subpopulations display functionally counterbalanced proportions in the DMS

In light of the failed attempts to restore goal-direction in aged mice through manipulations of the conditions of learning, we explored whether functional signatures of the aged striatum could inform about its incapacity to integrate the A→O contingencies necessary for goal-directed control. Recent findings in our lab highlight the importance of the territorial rules established by the two main neuronal subpopulations of the striatum (D1 and D2-SPNs), where activated D2-SPNs appear to exert regulatory influence over neighbouring D1-SPNs to influence outdated goal-directed learning in the DMS (Matamales et al., 2020). Here, to evaluate the status of this functional binary mosaic in the aged striatum, we mapped and classified large numbers of transcriptionally active SPNs in whole striatal sections of young and aged *drd2*-eGFP mice performing instrumental actions (Matamales et al., 2016, 2017, 2020) (Extended Data Fig. 7). We exposed both age groups to 3 extra days of A→O training (Fig. 4a and b) and evaluated the transcriptional state of striatal subpopulations in the DMS through detection of nucleosomal responses accumulated in D1- and D2-SPNs after 20 minutes of instrumental performance (day 17, Fig. 4c-e). We found that transcriptionally active (ta) D1- and D2-SPNs distributed similarly throughout medial and lateral territories of the dorsal striatum in young and aged mice, with abundant spread of both subpopulations in medial territories and a strong presence of taD1-but not taD2-SPNs in lateral areas, following the distribution patterns observed in earlier studies (Matamales et al., 2020). Despite equivalent general distribution patterns across age groups, the overall population of activated nuclei in each neuronal type was much reduced in aged as compared to young mice (Fig. 4c). We compared the levels of taD1- and taD2-SPNs with those collected in non-contingent control mice (A||O) exposed to the same training sessions but where the delivery of outcomes was bound to the actions performed by the other behavioural groups, and so remained independent of their lever press performance. Both A||O and aged contingent mice displayed similar activation levels in both subpopulations, which were significantly lower than those collected in young contingent mice (Fig. 4d, Supplementary Table 1), implying that A→O learning encoding was compromised in the aged group. Additional spatial analysis of taD1- and ta-D2-SPN distributions revealed that while young striata displayed a disproportionate number of taD1-SPNs that was propitious for plasticity in the DMS (reflected taD2/taD1-SPN ratios <1, Fig. 4e), the aged DMS maintained equal proportions of both neuronal types (taD2/taD1∼1, Fig. 4f), suggesting that plasticity in DMS D1-SPNs could be kept in check due to a functionally counterbalanced mosaic in the aged, similar to the D2-to D1-SPN regulatory process observed in extinction learning (Matamales et al., 2020).

**Fig. 4.**
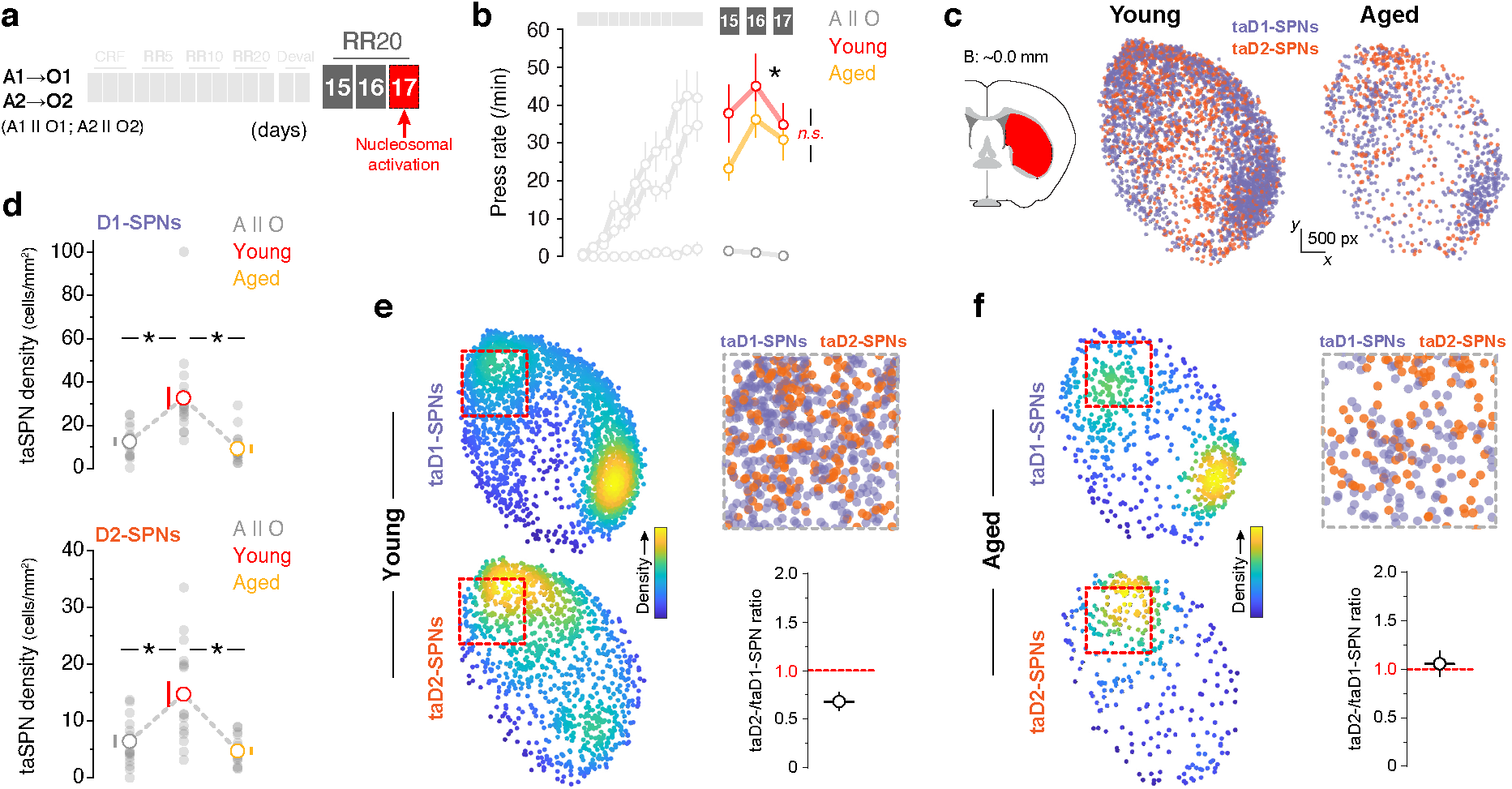
Instrumental action in aged mice is supported by a weak counterbalanced plasticity in striatal projection systems. **a**, After A1→O1 and A2→O2 training under increasing random ratio (RR) schedules (Fig. 2a) and outcome-specific devaluation tests (Deval; Fig. 3a), *drd2*-eGFP mice were re-trained on RR20 for 3 days prior to analysis of nucleosomal activation. A group with noncontingent A→O training (A1||O1; A2||O2) was added as a control (A||O control). **b**, Lever presses per minute (press rate) displayed by each group throughout instrumental re-training (averaged A1 and A2; 8 mice per group). **c**, Distribution of taD1- and taD2-SPNs in the posterior striatum (left) of all young and aged animals trained in (b). Scatter plots show the raw distribution of transcriptionally-active neurons in both populations superimposed. **d**, Densities of transcriptionally-active (ta) D1-SPNs (top) and D2-SPNs (bottom) in the striatum of mice trained in b. Light grey dots are individual hemisections. **e**-**f**, Scatter plots showing separate spatial distributions of taD1- and taD2-SPNs with pseudocoloured densities highlighting clustering (stacked hemisections). Insets show zoomed taD1- and taD2-SPN distributions in the posterior dorsomedial striatum (pDMS; top-right) and the corresponding taD2/taD1-SPN ratios calculated in the pDMS (red line denotes one-to-one; bottom-right). Data are mean ± SEM; 16 hemisections, 8 animals per group; *, overall/simple effect (black) and interaction (red). *n*.*s*., not significant (Supplementary Table 1).

### Mimicking young D2-to D1-SPN functional correspondence in the aged DMS enables vigorous goal-direction

Based on these findings and considering the functional logic of the striatal binary mosaic, we hypothesised that the sustained levels of D2-to D1-SPN transmodulation resulting from an aged, counterbalanced striatum could be keeping plasticity silent in DMS D1-SPNs. Such plasticity-lean state could prevent the addition of learning in medial basal ganglia systems that is necessary for the encoding of goal-direction (Balleine, 2019)—an effect that could prompt an early bias towards autonomous behavioural control (Yin et al., 2005). We sought a way to reintroduce in aged mice a long-lasting functional state that breaks the functional stillness of the striatal mosaic so that competence for A→O encoding returns, taking as a reference the plasticity status found in the young DMS (cf. Fig. 4e). We identified two factors that were critical for achieving a meaningful gain-of-function. First, the manipulation had to be durable and promoted early in learning, so that the desired plasticity state is achieved and self-maintained during the periods of A→O encoding. Second, endogenous plasticity occurring in D1-SPNs could not be enhanced non-specifically—any form of direct artificial functional supplement would add noise to the intrinsic cellular state that represents the A→O contingency, therefore diluting (rather than strengthening) essential D1-SPN function (Balleine et al., 2021).

We reasoned that inducing an early molecular de-sensitisation in D2-SPNs of the DMS could comply with both requirements: the inflicted change would (1) be long-lasting and thus present during critical periods of A→O encoding, and (2) result in a higher representation of D1-SPN plasticity, based on the D2-to D1-neuron transmodulation principle (Matamales et al., 2020), thereby leaving the synaptic state associated with the A→O contingency intact. Capitalising on the well-known desensitisation effects of PKC signalling on a variety of G protein-coupled receptors (GPCRs) in neurons (Bailey et al., 2009; Déry et al., 2001; Gereau and Heinemann, 1998; Namkung and Sibley, 2004), we sought to artificially enhance PKC signalling in DMS D2-SPNs of aged mice while undergoing instrumental learning. We expressed the synthetic hM3Dq-DREADD GPCR, a potent inductor of Gαq/11 signalling associated with PKC activation (Roth, 2016), selectively in D2-SPNs of the DMS of aged *adora2a*-Cre::*drd2*-eGFP hybrid mice (hereafter referred to as D2-hM3Dq-eGFP mice, Fig. 5a). We confirmed the induction of the Gαq/11/PLC/PKC/MAPK pathway (Trakul and Rosner, 2005) selectively in hM3Dq-transduced D2-SPNs 30 minutes after CNO administration (Fig. 5b, Extended Data Fig. 8a-f). This activation did not cross-react with cAMP signalling, since both nucleosomal and ribosomal responses dependent on cAMP signalling in D2-SPNs (Bertran-Gonzalez et al., 2009; Valjent et al., 2011) remained silent (Extended Data Fig. 8g and h).

**Fig. 5.**
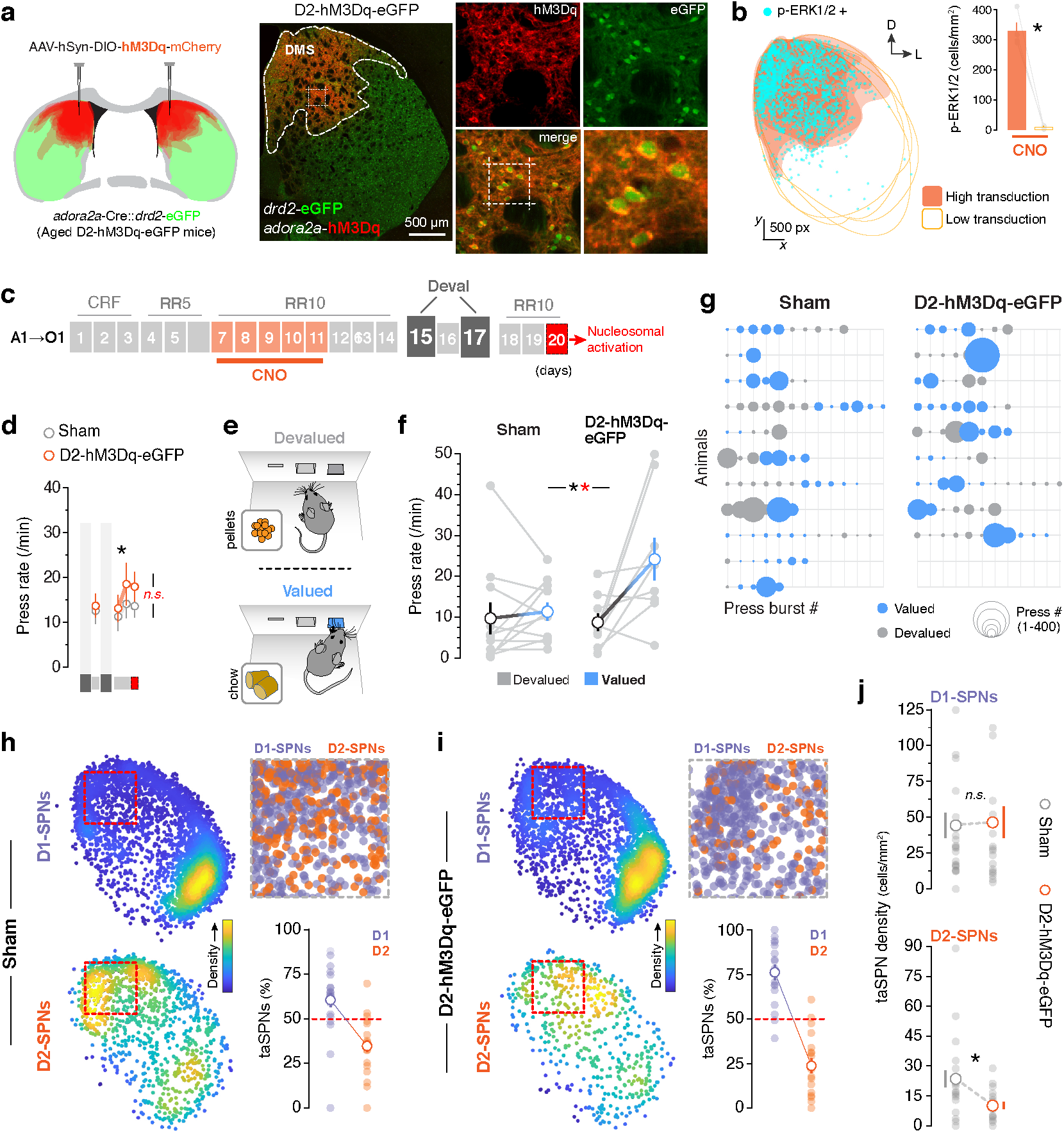
Long-lasting adjustment of D2-SPN plasticity during A→O encoding restores goal-direction in aged mice. **a**, An AAV expressing Cre-dependent hM3Dq-mCherry fusion construct or its vehicle was transduced selectively in pDMS D2-SPNs of aged *adora2a*-Cre::*drd2*-eGFP hybrid mice (D2-hM3Dq-eGFP and Sham). Left diagram shows the viral spread in all hemisections superimposed. Right panels: low and high magnification confocal images showing the selectivity of hM3Dq-mCherry expression based on hM3Dq – eGFP colocalisation. **b**, Characterisation of the chemogenetic effect detected 30 minutes after CNO injection. Left: mapping of activated (phospho-ERK1/2(+)) neurons across the striatum in superimposed hemisections. Orange territories denote areas with elevated viral transduction. Right: quantification of p-ERK1/2+ neuron density in high and low viral transduction areas. See Extended Data Fig. 8. **c**, Mice were trained on a single action (A1) → outcome (O1) contingency under continuous reinforcement (CRF) followed by increasing random ratio (RR) reinforcement schedules for 14 days. CNO (3 mg/kg, i.p.) was injected on days 7-11 30 mins prior to training (see Extended Data Fig. 9). On days 15 and 17, animals were exposed to outcome devaluation and test, with one re-training day in-between. The mice were then re-trained on RR10 for 3 days prior to analysis of nucleosomal activation. **d**, Lever presses per minute (press rate) displayed by each group throughout instrumental re-training. **e**, During outcome-devaluation procedure, mice were sated for 1 hr on either grain pellets (previously earned instrumentally; outcome is devalued) or regular chow food (outcome remains valued) prior to a lever pressing test. **f**, Press rates on the lever after satiation to pellets (devalued, grey) or after satiation to chow (valued, blue). Pairs of data points (light grey) are individual mice in devalued and valued trials. **g**, Bubble plot representing the response vigour exerted on the lever after pellet (devalued, grey) or chow (valued, blue) satiety throughout the lever pressing test. Press bursts are defined as all performance within a 20 second interval, and are organised chronologically. **h**-**i**, Distribution of taD1- and taD2-SPNs in the posterior striatum of all Sham (g) and D2-hM3Dq (e) animals after RR10 re-training. Scatter plots (left) show separate spatial distributions of taD1- and taD2-SPNs with pseudocoloured densities highlighting clustering (stacked hemisections). Inset (top-right) shows zoomed taD1- and taD2-SPN distributions in the pDMS. Bottom-right graphs show the taD2/taD1-SPN ratio calculated in the pDMS (red line denotes one-to-one). **j**, Densities of transcriptionally-active (ta) D1-SPNs (top) and D2-SPNs (bottom) in the pDMS. Light grey dots are individual hemisections. Data are mean ± SEM; 18-22 hemisections in 9-11 mice; *, overall/simple effect (black) and interaction (red). *n*.*s*., not significant (Supplementary Table 1).

We exposed aged D2-hM3Dq-eGFP mice and their Sham controls to the simplest form of instrumental learning where only one A→O contingency was trained, following the procedures used in Fig. 1a, with the aim of reducing conditions of learning confounds associated with task complexity and choice (Kosaki and Dickinson, 2010) (e.g. Figs 2 and 3). We administered CNO (3 mg/kg, i.p.) to both groups 30 mins prior to the training session for 5 consecutive days starting on day 7—point at which, in our first experiment, young and aged lever press performance had not yet started to diverge (Fig. 5c, cf. Fig 1b). This treatment did not affect overall instrumental performance, which increased equally in both D2-hM3Dq-eGFP and Sham groups as training progressed (Extended Data Fig. 9a and b, Fig. 5d, Supplementary Table 1). Unexpectedly, in contrast to our first experiment, both aged transgenic groups developed concatenated action sequences throughout instrumental training (Extended Data Fig. 9c-f), an effect that could be due to the improved behavioural performance in F1 hybrid strains (Sloin et al., 2022), supporting the notion that compromised action structure in ageing is a labile state that can be relatively easy to rescue through indirect interventions (Matamales et al., 2017) (e.g. Fig. 2). We next assessed whether the chemogenetic manipulation during training prompted goal-directed control by exposing both D2-hM3Dq-eGFP and Sham aged groups to single outcome devaluation procedures (Fisher et al., 2020): on days 15 and 17, mice were sated on either the instrumentally-conditioned outcome (devalued O) or regular chow (valued O) for 1h followed by an extinction test, in which the trained lever was present but no outcome was delivered (Fig. 5e). We found that while Sham animals displayed similar lever press rates across tests, D2-hM3Dq-GFP mice showed significantly elevated lever press rates in valued vs. devalued extinction tests (Fig. 5f, Supplementary Table 1). This effect was supported by a more vigorous approach to the lever recorded in D2-hM3Dq-GFP mice but not Sham controls during valued vs. devalued tests (Fig. 5g).

We next re-trained both groups on the same A→O contingency for 3 days with the aim of confirming that our chemogenetic desensitisation conducted early in training had resulted in long-lasting decompensation of D1- and D2-SPN plasticity nearly 10 days after treatment ceased. Unlike the aged group in our initial experiment (cf. Fig. 4f), we found that Sham animals displayed a certain level of plasticity imbalance with a slightly higher proportion of taD1-than taD2-SPNs in the DMS (Fig 5h) although this taD1/taD2-SPN disproportion was greatly pronounced in D2-hM3Dq-GFP mice (Fig. 5i). Importantly, this disproportion in D2-hM3Dq-eGFP mice was due to a specific reduction of taD2-SPN density and not an increase in taD1-SPN numbers (Fig. 5j, Supplementary Table 1). The fact that both groups retained comparable densities of taD1-SPNs suggests that our manipulation did not interfere with the essential cellular state associated with the integration of the A→O contingency, which is proposed to be encoded by D1-SPNs (Balleine et al., 2021). These data suggest that an adequate state of functional D2-to D1-SPN correspondence is required to render the striatal mosaic competent for A→O integration, and that manipulations approximating such a state can unleash capacity for goal-directed control in the maladjusted DMS.

## Discussion

In this study we sought to determine the origin of the propensity of aged animals to express autonomous stimulus→response (S→R) control of behaviour by disambiguating the influence of factors relating to the *conditions of learning* from those relating to the *state of the brain*. We found that instrumental performance was clearly degraded in aged mice, something that could a priori account for altered conditions of learning that hinder capacity to goal-directed control. However, while such alterations in action structure were easily recovered by introducing elements of task complexity such as training to additional action→outcome (A→O) contingencies in various ways, none of these manipulations were effective at enabling goal-direction in aged mice. Instead, we found that aged mice conducting instrumental behaviour presented unusual one-to-one proportions of transcriptionally-active (ta) D1- and D2-SPNs in the dorsomedial striatum (DMS). We hypothesised that this functional counterbalance could promote a state of plasticity quietness that prevented the appropriate integration of A→O learning, based on our recent studies highlighting the critical role of local modulatory interactions amongst SPN subpopulations in defining striatal function (Matamales et al., 2020). We showed in aged transgenic mice that introducing biases to the functional D2-to D1-SPN correspondence through cell-specific chemogenetics in the DMS restored the animals’ ability to integrate A→ O contingencies, something that allowed the expression of vigorous, cognitively guided, goal-directed action.

The lack of sequential action structure reported here in aged mice has been observed in humans and could, by itself, explain a deficient encoding of the A→O contingency based on rate correlation views: as the ratio requirement increases, a commensurate extension of the action sequences would appear necessary to provide adequate feedback and maintain a proportional A→O correspondence (Dickinson, 1985; Perez and Dickinson, 2020). To the animals’ perception, sequences truncated by premature magazine checks introduce unsuccessful performance and, with it, deviations from the A→O linearity, which can very well compromise the integrity of the contingency during learning. In support of this, training on interval schedules of reinforcement—known to promote poor lever press performance (Dickinson et al., 1983) as well as low A→O correspondence (Dickinson, 1985; Perez and Dickinson, 2020)—often lead to autonomous behaviour relatively easily, something that can be preven-ted, for example, if training on a second A→O contingency is introduced (Kosaki and Dickinson, 2010). In our hands, interval schedules also produced poor lever press performance in both young and aged mice, and introducing a second contingency was insufficient to promote goal-direction in any mouse, regardless of age. While these results failed to reproduce the effects by Kosaki and Dickinson in rats, they still support the idea that maintaining a good A→ O correlation may be essential for acquiring and/or maintaining goal-direction. However, our next set of experiments showed that restoring performance (and with it, A→O linearity) was not sufficient to secure goal-direction in aged mice, suggesting that factors other than the behavioural conditions during instrumental learning were responsible for deficient A→O encoding in ageing.

Accordingly, we found in aged mice conducting instrumental conditioning tasks an abnormal pattern of striatal engagement that was characterised by an equal confluence of transcriptionally active (ta) striatal projection neuron (SPN) subpopulations in the DMS. Such pattern could be a consequence of the known functional declines commonly observed in aged fronto-striatal networks (Eppinger et al., 2011), or could be even related to the compensatory overactivations seen in prefrontal and fronto-striatal circuits in aged brains (Matamales et al., 2017; Park and Reuter-Lorenz, 2009; Zhang et al., 2017). Regardless, the abnormal corticostriatal drive in fronto-medial regions could, in time, synchronise molecular activities across both D1- and D2-SPN subpopulations, bringing up a state of plasticity stillness in the medial striatum. This process follows the logic of the functional binary mosaic, a recently described view of striatal function that highlights the importance of the neuromodulatory rules established locally by D1- and D2-SPNs, such as the D2-to D1-SPN transmodulation. This model relies on the large molecular capacities of SPNs (Greengard, 2001) and is useful to define the functional territoriality that supports the integration of learning in the striatal continuum (Matamales et al., 2020). We found in this study that aged brains display a proportionate molecular engagement of both D1- and D2-SPNs in the DMS, which, we propose, reflects a state of counterbalanced plasticity by which D1-SPNs cannot break free from the close to one-to-one modulation exerted by neighbouring D2-SPNs. Such counterbalanced state is similar to the regulatory D2-to D1-SPN transmodulation described in extinction learning, where previously acquired contingencies encoded in DMS D1-SPNs seem to be re-programmed by D2-SPNs after these functionally colonise the same striatal space (Matamales et al., 2020). In line with this, we show here that breaking this static functional balance by long-lastingly reducing the proportion of transcriptionally active D2-SPNs proved sufficient to restore plasticity capacity in the aged DMS, as reflected by the gained ability of these mice to integrate A→O contingencies and behave goal-directedly.

The main result in this study involves a disrupted ability to encode A→O contingencies in aged mice. We here replicated this result in four different experimental settings: (1) when two parallel A→O relationships were trained sequentially (e.g. Fig. 3c); (2) when two parallel A→O relationships were trained interspersedly (Fig. 3F); (3) when two parallel A→O relationships were trained under interval schedules (e.g. Ext Data Fig. 6); and when one single A→O relationship was trained on its own (Fig. 5). While this resilient behavioural feature is indicative of a defective encoding of A→O contingencies necessary for goal-directed action in ageing and is in line with previous literature, we do not intend to claim that all aged brains will present faulty A→O encoding, or will do so at this same stage of learning. For example, in a previous study in a different animal facility and using a different strain of mice (Matamales et al., 2016), we found that A→O encoding was sufficient initially, but was faulty in subsequent learning when contingencies were reversed, an effect that also related to medial striatal dysfunction. Brain ageing is a complex multifactorial process that involves various forms of functional decline and compensation, and as such its biological features in the general population are variable and likely determined by tendencies and trends rather than discrete processes.

Our results a priori support the views that aged brains show a higher reliance on anteced-ent S→R processes and may consequently show enhanced propensity to autonomous behavioural control (Eppinger et al., 2013; Ito et al., 2021; Mata et al., 2010; de Wit et al., 2014). However, such conclusion should be considered carefully—whether propensity to S→ R-like behaviour reflects a core recruitment of a “habit”, model-free, controller as a feature of the aged brain or is simply a consequence of a faulty integration of A→O encoding at any point of learning remains to be answered (Balleine and Dezfouli, 2019). Our experiments in mice do not allow to directly disambiguate this question, although they support the notion that such propensity to autonomous responding may originate from an identifiable neuronal pattern that keeps goal-directed control dormant. Collectively, the specific features of striatal function in ageing reported here, as well as the neural strategies proposed to restore them, provide critical information about how goal-directed and autonomous controllers may be represented in striatal circuits (O’Hare et al., 2016), and open the door to interventions that help bringing back the dominance of cognitive control of behaviour in autonomy-prone brains—such as the aged brain.

## Acknowledgements

We thank Jennifer Strempel, Anne Rowan and Lydia Williams for assistance with animal care.

