## Supplementary Materials for "Restoring functional D2- to D1-neuron correspondence enables goal-directed action control in long-lived striatal circuits"

www.NeuroModuLab.org

**This PDF file includes:**

Supplementary Methods
Supplementary Tables
Extended Data Figures 1-9

#### Supplementary Methods

(references in main article)

##### Animals

All experimental procedures were approved by the Animal Care and Ethics Committee at UNSW Sydney in accordance with the Australian Code of Practice for the Care and Use of Animals for Scientific Purposes (National Health and Medical Research Council, 2013). Adult mice (2-6 months old in Young groups and 18-22 months old in Aged groups) of both genders were used in these experiments. Behavioural experiments were performed either on C57BL6/J mice (RRID:IMSR_JAX:000664) purchased from the Animal Resources Centre (Perth, Australia) or outbred *drd2*-eGFP with either C57BL6/J (C57BL6/J x *drd2*-eGFP$^{+/-}$, here referred to as *drd2*-eGFP) or with C57Bl6/J-backcrossed *adora2a*-Cre$^{+/+}$ (*adora2a*-Cre$^{+/-}$ x *drd2*-eGFP$^{+/-}$ F1 hybrids, here referred to as *adora2a*-Cre::*drd2*-eGFP). The original transgenic mouse lines were purchased from the Mutant Mouse Resource and Research Centres (MMRRC, NIH). The *drd2*-eGFP line (B6(Cg)-Tg(Drd2-EGFP)S118Gsat/KreMmucd [RRID:MMRRC_036931-UCD]) expresses eGFP in D2-SPNs. The *adora2a*-Cre line (STOCK Tg(Adora2a-cre)KG139Gsat/Mmucd [RRID:MMRRC_031168-UCD]) expresses Cre in D2-SPNs. Mice were housed in individually ventilated plastic home cages with a stand-alone air handling unit (Tecniplast, IT) in a climate-controlled colony room on a 12-h light/dark cycle, and were allowed ad libitum access to water. Littermates in each home cage were maintained throughout the experiment, and were randomly allocated to each experimental group.

##### Behavioural procedures and analysis

*Apparatus.* All training procedures were carried out in sound- and light-resistant modular operant chambers (Med Associates Inc, PO). The chambers were illuminated with a 3 W, 24 V house light. Each chamber contained a recessed feeding magazine at the centre of the right-side wall connected to an individual food dispenser that could deliver either 20 mg grain-based (#F0163) or 20 mg purified (#F0071) Dustless Precision Pellets (Bio-Serv, NJ) into the magazine when activated. Either side of the magazine contained retractable levers. Med-PC software was used to direct the insertion and retraction of the levers, illumination of the light and delivery of the pellets. The software allowed the recording of timestamp events each 1 msec, including the number of lever presses, magazine entries and food pellets delivered throughout the duration of the experiment. A combination of Med State Notation language and customized MATLAB (MathWorks, Natick, MA) scripts were used to retrieve timestamp data. Activity was also monitored using D-Link DCS-932L IP LED infrared cameras (VGA 1/5 inch CMOS sensor with built-in microphone) through D-ViewCam Software (D-link Corporation, Taiwan).

*Magazine training.* Food intake was restricted throughout the experiments, with mice being maintained at approximately 85% of their free-feeding weight. Prior to all instrumental procedures, mice were exposed to three to four days of magazine training in the operant chambers (1 session/day). Each mouse was assigned one operant chamber, which was maintained throughout the duration of the experiment. The chamber light turned on to signal the beginning of the session and off at its termination. Both levers were retracted and mice explored the chamber freely. Twenty grain-based food pellets (in experiments involving one lever → outcome training) or forty pellets (20 grain-based and 20 purified in experiments involving two lever → outcome contingencies) were delivered into the magazine at random time intervals (RT 60 seconds) over ~30 and ~60 min, respectively. Consumption of pellets was monitored at the end of each session.

*Single lever instrumental training.* These procedures were based on one lever → outcome operant conditioning, where an action (lever press) was associated with the delivery of an outcome (20 mg grain-based pellet). Mice were trained once daily. Each training session began when the test chamber light turned on and the lever contingent to the delivery of the pellets inserted in the chamber, and ended when the chamber light turned off and the lever retracted. Assignment of right or left lever was counterbalanced across mice. Each training session continued until mice had received a total of 20 pellets, or the session timed out at 30 min. For the first 3 days of instrumental training (days 1-3), the lever delivered the pellet on a continuous reinforcement (CRF) schedule, in which every lever press was rewarded. This was followed by a random ratio (RR) reinforcement schedule, in which the probability of the outcome given a lever response was first reduced to 0.2 (RR5; days 4-5) and then to 0.1 (RR10; days 7-15 [Fig. 1] or 7-14 [Fig.5]).

*Two-lever instrumental conditioning (sequential).* In most two-lever experiments, two action – outcome contingencies were trained in parallel but separately. For each action → outcome contingency, training session began with the illumination of the chamber light and insertion of one lever into the operant box, and ended with the extinguishing of the chamber light and lever retraction. The mice were exposed to two consecutive training sessions each day where only one lever was inserted into the box (and only the corresponding pellet delivered). The order of the sessions (purified or grain pellet first) was randomly alternated each day and lever-outcome assignment was fully counterbalanced. Pressing of the lever (right or left; A1 or A2) resulted in the delivery of the associated pellet (grain or purified; O1 or O2). Each training session (A1→O1 and A2→O2) continued until mice had received a total of 20 pellets, or the session timed out at 30 min. For the first three days of instrumental training (days 1-3), mice were trained on a constant reinforcement schedule (CRF); every lever pressing action was rewarded with the corresponding outcome. The probability of the outcome given a response (P) was gradually shifted over the following days of training using increasing random ratio (RR) schedules of reinforcement: a RR5 schedule (P = 0.2) was used on days 4–6, RR10 (P = 0.1) was used on days 7–9 and RR20 (P = 0.05) was used on days 10-12.

*Two-lever instrumental conditioning (interspersed).* On a subset of experiments, the two action – outcome contingencies were trained in parallel but in interspersed trials within the same session. Training sessions began with the illumination of the chamber light and insertion of one lever (left or right, picked randomly) into the operant box. In each trial, animals were required to earn 5 of the corresponding rewards at the required schedule for it to end, at which point the lever retracted and a pseudorandom inter trial interval (ITI; 45 seconds in average) elapsed before the onset of the next trial. The session ended either when 40 rewards (of whatever kind) had been earned, or when 60 min had elapsed (whichever came first), at which point the chamber light extinguished and the lever retracted. Schedules of reinforcement and the days at which they were trained were the same as the sequential version of the task (above).

*Two-lever instrumental conditioning (interval schedules).* On another subset of experiments, the two action – outcome contingencies were trained in parallel and in two separate sessions each day (as above), but using interval instead of ratio schedules of reinforcement. On these, each training session (A1→O1 and A2→O2) continued until mice had received a total of 40 pellets, or the session timed out at 60 min. For the first three days of instrumental training (days 1-3), mice were trained on a constant reinforcement schedule (CRF); every lever pressing action was rewarded with the corresponding outcome. On the following days, lever pressing was rendered contingent to outcome delivery under increasing random interval (RI) schedules of reinforcement. On days 4-6, periods of contingency during the session were defined on every second by P = 0.03 (RI30), whereas on days 7-17 probability was shifted to P = 0.016 (RI60).

*Devaluation of outcomes and contingency test.* On the last three days of initial instrumental training, mice were pre-habituated to devaluation boxes for 30 min to prevent novelty-induced stress on the days of devaluation. Pre-habituation was performed in a different room, where mice were placed individually into squared boxes which contained an empty pellet container. In single-lever experiments, two devaluation trials were performed, starting the day after the last day of training and including a re-training session in-between (days 15 and 17). On each devaluation day, mice were exposed to ~50 g of either (instrumentally-earned) grain pellets or regular chow for 1h (counterbalanced) followed by a 10-min extinction test aimed at evaluating the state of the A→O contingency. In such test, each mouse was placed in their assigned operant chambers, and the relevant lever was inserted but no outcome was delivered. In two-lever experiments, the two devaluation trials were consecutive (days 13 and 14), with outcome being devalued each day. The pellet container was filled with approximately 50 g of one (e.g. grain pellets) or another (e.g. purified pellets) outcome and animals were allowed to consume them to satiety over a 1 hr exposure period. Immediately following exposure, animals underwent an outcome-specific devaluation choice test in their assigned operant chambers, where the two levers where inserted but no outcomes were delivered. In both single- and double-lever experiments, the day after the last test, mice were re-trained on the A→O contingencies and schedule they had immediately before devaluation and test for another 3 days, prior to neuronal activation analyses.

*Noncontingent instrumental training.* In neuronal mapping experiments (two-lever training, RR schedules), a noncontingent group (A1||O1, A2||O2; 8 mice) was trained in parallel to control for any neuronal activation that was possibly due to exposure to any associative process between the context and the reinforcer except for the A→O contingency. These mice received a reward whenever their ‘contingent’ littermate earned it through lever pressing;

they were exposed to the same conditions and food intake as their instrumental directors, but did not develop any action-outcome contingency as reward delivery was to another mouse's actions.

*Analysis and visualisation of behavioural data.* Individual events (left or right lever press, magazine entries, and pellet delivery) that occurred during each training session were recorded. The time at which each event occurred was stamped with a 10 ms resolution. Datasets were imported into MATLAB and different approaches were used to analyse and visualise the data. Discrete sequences and their initiation, execution and termination elements were identified based on the interspacing of lever presses and magazine checks. Consecutive magazine checks produced after the termination of a sequence were ignored, as they were the result of the consumption of reward. Parameters such as concatenation (number of presses per sequence) and sequence duration (time elapsed from initiation to termination) were calculated at this point for each sequence, and average values were obtained for each mouse. To represent the automatisation of instrumental action throughout training we plotted return maps of inter press interval (IPI) collected in each animal for a given training session. Each graph shows the stacked maps all mice from a particular group on the indicated day of training. Each dot is an IPI and includes the time delay to its preceding (x) and succeeding (y) element in the sequence. The colour for each data point in the heatmap scale is based on their calculated kernel density estimates (Botev et al., 2010). Data are Log10 transformed to approximate values. To visualise response concatenation, all events corresponding to the delivery of pellets throughout training were sorted chronologically, and the data points were colour coded according to the number of lever presses leading to each reward. To represent the response vigour on each lever during devaluation test, we defined a response cluster as bursts of lever presses separated by intervals of at least 20 seconds. We plotted the identified bursts chronologically using a bubble chart where the size of single data points is based on the number of lever presses within the cluster.

**Viral procedures**

*Inoculation of AAVs.* Live viruses were injected into the brain through stereotaxic surgery. Mice were pre-anesthetised in an induction chamber with oxygen/isoflurane mixture (Laser Animal Health, Pharmachem, Australia). Pedal reflex was used to monitor anaesthesia before placing the mouse in the stereotaxic frame (Kopf Instruments) fitted with a mask supplying a continuous flow of oxygen/isoflurane mixture (1L/min oxygen, 1.5 ml/min isoflurane). Once deeply anesthetised, a 1.5 cm incision was made to expose the skull. For each microinjection, a small hole was pierced (~0.2 mm width) at the appropriate R-C and M-L coordinates (see above). A glass capillary (3.5", Drummond Scientific #3-000-203-G/X) was pulled using a micropipette puller (P-97, Sutter Instrument), pre-filled with mineral oil fitted into a Nanoject III (Drummond Scientific). The AAV-loaded micropipette was slowly inserted vertically through each hole until reaching target (D-V coordinate from skull surface). Once in target and after a 2-min waiting interval, the solution was injected at a rate of 1 nl/sec for a total of 10 minutes. Five min after the end of the injection, the pipette was gently pulled out, and the incision was closed with surgery suture silk and sealed with tissue adhesive (3M Vetbond). Animals were placed in cages separately on a warming plate (37 ºC) during recovery. A pre-operative s.c. administration of metacam (5 mg/kg) ensured analgesia during and after the intervention. Behavioural procedures started after a 3-week recovery period. None of the mice displayed perceptible wounds by the start of the behavioural experiments.

*Chemogenetics.* Stimulation of molecular signalling activity specifically in D2-SPNs of the posterior dorsomedial striatum (pDMS) was achieved through bilateral stereotaxic microinjection of Cre-dependent hM3Dq DREADD virus (AAV-hSyn-DIO-hM4D(Gi)-mCherry, UNC Vector Core, NC) in aged *adora2a*-Cre::*drd2*-eGFP hybrid mice (coordinates: R-C: +0.13 mm [bregma]; M-L: 1.70 mm [bregma]; D-V: -2.65 mm [skull surface]) combined with an intraperitoneal injection of clozapine N-oxyde (CNO; dissolved in saline to 3 mg/kg; NIMH Chemical Synthesis and Drug Supply Program, Bethesda, MD). In order to validate the effects of this manipulation, particularly in aged animals, two of these mice received a single CNO injection prior to rapid intracardial perfusion and neuronal activation mapping. Animals used for two-lever instrumental conditioning were injected with CNO on the relevant days 30 min prior to the start of the training session.

**Tissue processing and immunofluorescence labelling**

*Transcardial fixation and tissue sectioning.* While still conducting instrumental behaviours (20 minutes in), mice were rapidly anesthetised by exposure to 10 sec of isoflurane gas in an induction chamber (4% in air; Laser Animal Health,

Pharmachem, Australia), followed by an intraperitoneal injection of sodium pentobarbital (0.5 mL of 10% Lethabarb; Virbac Pty. Ltd., Australia). After ensuring deep anaesthesia by testing paw and tail reflexes, mice were perfused transcardially using an air pressure system (constant flow of 15 ml/min) with cold paraformaldehyde solution (4%, w/v) in phosphate buffer saline (pH 7.4). Brains were dissected out and post-fixed overnight in 4% paraformaldehyde solution at 4°C. Consecutive 30 μm coronal sections spanning the rostro-caudal extent of the striatum were obtained for each animal using a vibratome (VT1000s, Leica Microsystems, Germany). Free-floating slices were stored at -20°C in an anti-freeze cryoprotectant solution (ethylene glycol, 30% v/v; glycerol, 30% v/v, 0.25 M Tris buffer) until processed for immunofluorescence.

*Immunofluorescence staining.* Free floating brain slices (bregma: ~0.14 mm) were rinsed 3 times in Tris-buffered saline solution (TBS, 0.5 M) for 10 min on an orbital shaker at room temperature (R.T.) prior to membrane permeabilization treatment with triton X-100 solution (0.2%, v/v in TBS), which was applied for 1 h 30 min at R.T. After 3 10-min washes with TBS, slices were incubated at 4°C on an orbital shaker with rabbit polyclonal anti-phospho Histone H3 (Ser10) antibodies (p-H3; 1:500 dilution; Millipore, Cat. No. 06-570), polyclonal anti phospho-ERK1/2 (p-ERK1/2, 1:300 dilution, Cell Signalling Technology Cat. No. 9102) or anti-phosphorylated ribosomal protein S6 (p-rpS6; 1:300; Cell Signaling Technology, #2215). Following a 48-h incubation period, unbound primary antibodies were washed off with 4 washes of TBS, and bound primary antibodies were detected through incubation for 1 h 30 min at room temperature with donkey anti-rabbit Alexa Fluor 546 secondary antibodies (1:400 dilution; Thermo Fisher Scientific; Cat. No. A10040). Unbound antibodies were washed off through 2 10-min rinses in TBS and 2 10-min final washes in TB. Brain samples were then placed on glass slides and mounted with a coverslip on Vectashield antifade mounting medium (Vector Laboratories, CA; Cat. No. H-1000). Slides were stored at 4°C in the dark until image acquisition.

**Image acquisition and quantitative analysis**

*Spinning disk confocal microscopy.* Fluorescently-labelled brain sections were imaged using a confocal field scanning spinning disk system equipped with the Diskovery multi-modal imaging platform (Andor Technology) and a Zyla 4.2 sCMOS camera (Andor Technology), that allows fast capture of large mosaic images and high-sensitivity high-dynamic range imaging. The Diskovery platform was added to a Nikon Eclipse TiE microscope body with a motorized stage and the Nikon Perfect Focus System, and image acquisition was controlled by Nikon NIS-Elements software. Single high-magnification wide field-of-view images for each brain hemisphere were generated by automatically stitching multiple adjacent frames (20X objective, 2x2 binning) from within a defined area using the motorized stage. Individual images were automatically aligned (registered) to a common reference image using StackRegJ plugin in ImageJ/Fiji software (Schindelin et al., 2012), such that orientation similarity in striata from different animals was maximal.

*Nucleosomal response mapping.* High-resolution mosaics of brain samples obtained with NIS-Elements were quantitatively analysed using ImageJ/Fiji software (Schindelin et al., 2012). The outer boundary of the striatum or defined subregions (e.g., High/Low viral transduction areas) were manually delineated in each hemisection (freehand tool), the area of the region was measured ($mm^2$), and coordinates (x, y) of the points defining the line were obtained. To quantify the number of activated SPNs in *drd2*-eGFP mice and retrieve their location across the striatum, images were automatically thresholded based on P-H3 immunolabelling and individual nuclei were automatically detected using the "Analyze Particle" command in Fiji, which finds the edge of an object (i.e. nucleus), outlines its contour using the wand tool, creates an individual region of interest (ROI), and then determines the coordinates (x, y) of its centre point (centroid). To determine whether an activated SPN was D1- or D2-type, the generated ROIs were overlaid on the eGFP image and the mean grey value (m.g.v) of all the pixels within the selection was then measured. Resulting neuronal data, including position (x, y coordinates), area and mean grey value of the eGFP channel for each detected ROI, were exported in a tab-delimited spreadsheet format and imported into MATLAB. Custom-made scripts were used to obtain the density of activated SPNs (P-H3 counts/$mm^2$) for each SPN population, based on an eGFP m.g.v threshold, which was obtained by averaging the eGFP m.g.v of a number of identified D1- and D2-SPNs. The position of activated neurons within the tissue was digitally reconstructed by combining a line plot defined by the edge points of the striatum and a scatter plot with circles at the locations specified by the neurons' centroid coordinates, color-coded according to their eGFP m.g.v. To visualise the density of activated neurons for each neural population, we colour coded single data points based on their calculated kernel density estimate (Botev et al., 2010). To calculate the density of neurons in defined subregions, we used the "inpolygon" function MATLAB, that returns the points that are inside the edge of the polygon area defined by the (x, y) coordinates of the ROI line.

**Statistical analyses**

Statistical significance of the effects found in our data was assessed using IBM SPSS Statistics software (versions 25-28; IBM Corporation, Somers, NY). The a priori alpha level was set at $p < .05$ (two-sided). To analyse the effect of one or more independent variables, Levene's test was first used to test the null hypothesis that the error variance of each dependent variable was equal across groups, and uni- or multi-factorial ANOVA was then conducted. If one or more repeatedly measured variables were considered, uni- or multifactorial repeated measures ANOVA was conducted. Homogeneity of variance was tested through Mauchly's test of sphericity, and Greenhouse-Geisser corrections were considered when sphericity was not met. Mixed ANOVA analyses were conducted in experiments involving within-subject and between subjects factors. In instances where further statistical detail was appropriate, we conducted additional simple effects comparisons based on individual one-way ANOVAs or Bonferroni post-hoc comparisons. Statistical comparisons between two means were conducted through independent t-tests, and equal variances were assumed or not according to the Levene's test for equality of variances.

#### Supplementary Tables

**Supplementary Table 1 | Statistical analyses.** Factor interactions are marked in red. ED = Extended Data Figures.

| Fig. | Analysis | Test | Dep. variable | Factor(s) | Statistic | P value |
|---|---|---|---|---|---|---|
| 1b | Instrumental acquisition | Two Way *mixed* ANOVA | Lever press rate (press/min) | (a) Training day (within-subjects)<br><br>(b) Group (between subjects) | Main Effect (Training): $F_{(14,196)} = 116.07$<br><br>Mauchly's sph. test: $p < 0.001$ Greenhouse Geisser correction: $F_{(3.22,45.1)} = 116.07$<br><br>Training x Group interaction: $F_{(3.22,45.1)} = 3.87$ | **< 0.001**<br><br>**< 0.05** |
| 1c | Presses/sequence | Two Way *mixed* ANOVA | Presses/sequence | (a) Day (within-subjects)<br><br>(b) Group (between subjects) | Main Effect (Day): $F_{(1, 14)} = 50.51$<br><br>Day x Group interaction: $F_{(1, 14)} = 32.13$ | **< 0.001**<br><br>**< 0.001** |
| 1c | Presses/sequence (young) | One Way *repeated measures* ANOVA | Presses/sequence | Day (within subjects) | $F_{(1, 7)} = 55.37$ | **< 0.001** |
| 1c | Presses/sequence (aged) | One Way *repeated measures* ANOVA | Presses/sequence | Day (within subjects) | $F_{(1, 7)} = 1.96$ | = 0.204 |
| 1d | Sequence duration | Two Way *mixed* ANOVA | Sequence duration (seconds) | (a) Day (within-subjects)<br><br>(b) Group (between subjects) | Main Effect (Day): $F_{(1, 14)} = 2.33$<br><br>Day x Group interaction: $F_{(1, 14)} = 11.84$ | = 0.149<br><br>**< 0.01** |
| 1d | Sequence duration (young) | One Way *repeated measures* ANOVA | Sequence duration (seconds) | Day (within subjects) | $F_{(1, 7)} = 9.19$ | **< 0.05** |
| 1d | Sequence duration (aged) | One Way *repeated measures* ANOVA | Sequence duration (seconds) | Day (within subjects) | $F_{(1, 7)} = 2.78$ | 0.14 |
| 1g | Concatenation | Two Way *mixed* ANOVA | Concatenated sequences (%) | (a) Day (within-subjects)<br><br>(b) Group (between subjects) | Main Effect (Day): $F_{(1, 14)} = 19.86$<br><br>Day x Group interaction: $F_{(1, 14)} = 5.46$ | **= 0.001**<br><br>**< 0.05** |
| 1g | Concatenation (young) | One Way *repeated measures* ANOVA | Sequence duration (seconds) | Day (within subjects) | $F_{(1, 7)} = 31.38$ | **= 0.001** |
| 1g | Concatenation (aged) | One Way *repeated measures* ANOVA | Sequence duration (seconds) | Day (within subjects) | $F_{(1, 7)} = 1.78$ | = 0.224 |

| Fig. | Analysis | Test | Dep. variable | Factor(s) | Statistic | P value |
|---|---|---|---|---|---|---|
| 2b | Instrumental acquisition | Two Way *mixed* ANOVA | Lever press rate (press/min) | (a) Training day (within-subjects)<br><br>(b) Group (between subjects) | Main Effect (Training): $F_{(11,154)} = 41.272$<br><br>Mauchly's sph. test: $p < 0.001$ Greenhouse Geisser correction: $F_{(3.57,49.99)} = 41.272$<br><br>Training x Group interaction: $F_{(3.57,49.99)} = 1.209$ | **< 0.001**<br><br>= 0.318 |
| 2d | Sequence length | Two Way *mixed* ANOVA | Presses/sequence | (a) Day (within-subjects)<br><br>(b) Group (between subjects) | Main Effect (Day): $F_{(2, 60)} = 51.72$<br><br>Day x Group interaction: $F_{(2, 60)} = 3.09$ | **< 0.001**<br><br>**> 0.05** |
| 2d | Sequence length (young) | One Way *repeated measures* ANOVA | Presses/sequence | Day (within subjects) | Main Effect (Day): $F_{(2,30)} = 79.29$<br><br>Mauchly's sph. test: $p < 0.01$ Greenhouse Geisser correction: $F_{(1.26,18.98)} = 79.29$ | **< 0.001** |
| 2d | Sequence length (Aged) | One Way *repeated measures* ANOVA | Presses/sequence | Day (within subjects) | $F_{(2,30)} = 10.11$ | **< 0.001** |
| 2e | Sequence duration | Two Way *mixed* ANOVA | Sequence duration (seconds) | (a) Day (within-subjects)<br><br>(b) Group (between subjects) | Main Effect (Day): $F_{(2,60)} = 10.45$<br><br>Mauchly's sph. test: $p < 0.001$ Greenhouse Geisser correction: $F_{(1.31,39.40)} = 10.45$<br><br>Training x Group interaction: $F_{(1.31,39.40)} = 0.80$ | **= 0.001**<br><br>= 0.844 |
| 2e | Sequence duration (young) | One Way *repeated measures* ANOVA | Sequence duration (seconds) | Day (within subjects) | Main Effect (Day): $F_{(2,30)} = 38.22$<br><br>Mauchly's sph. test: $p < 0.001$ Greenhouse Geisser correction: $F_{(1.2,18)} = 38.22$ | **< 0.001** |
| 2e | Sequence duration (aged) | One Way *repeated measures* ANOVA | Sequence duration (seconds) | Day (within subjects) | Main Effect (Day): $F_{(2,30)} = 3.183$<br><br>Mauchly's sph. test: $p < 0.01$ Greenhouse Geisser correction: $F_{(1.32,19.76)} = 3.183$ | = 0.056<br><br>= 0.08 |
| 2g | Concatenation | Two Way *mixed* ANOVA | Concatenated sequences (%) | (a) Day (within-subjects)<br><br>(b) Group (between subjects) | Main Effect (Day): $F_{(2,60)} = 27.27$<br><br>Mauchly's sph. test: $p < 0.001$ Greenhouse Geisser correction: $F_{(1.42,61)} = 27.27$<br><br>Training x Group interaction: $F_{(1.42,61)} = 0.96$ | **< 0.001**<br><br>= 0.842 |

| Fig. | Analysis | Test | Dep. variable | Factor(s) | Statistic | P value |
|---|---|---|---|---|---|---|
| 2g | Concatenation (young) | One Way *repeated measures* ANOVA | Sequence duration (seconds) | Day (within subjects) | $F_{(2, 30)} = 13.04$ | **< 0.001** |
| 2g | Concatenation (aged) | One Way *repeated measures* ANOVA | Sequence duration (seconds) | Day (within subjects) | Main Effect (Day): $F_{(2,30)} = 14.57$ Mauchly's sph. test: $p < 0.01$ Greenhouse Geisser correction: $F_{(1.24,18.62)} = 14.57$ | **= 0.001** |
| ED 3b | Sequence length | Two Way *mixed* ANOVA | Presses/sequence | (a) Day (within-subjects) (b) Group (between subjects) | Main Effect (Day): $F_{(2, 52)} = 2.57$ Day x Group interaction: $F_{(2, 52)} = 2.89$ | = 0.086 = 0.064 |
| ED 3b | Sequence length (young) | One Way *repeated measures* ANOVA | Presses/sequence | Day (within subjects) | $F_{(2,22)} = 0.77$ | = 0.475 |
| ED 3b | Sequence length (aged) | One Way *repeated measures* ANOVA | Presses/sequence | Day (within subjects) | $F_{(2,30)} = 4.71$ | **< 0.05** |
| ED 3c | Sequence duration | Two Way *mixed* ANOVA | Sequence duration (seconds) | (a) Day (within-subjects) (b) Group (between subjects) | Main Effect (Day): $F_{(2, 52)} = 9.34$ Day x Group interaction: $F_{(2, 52)} = 0.284$ | **< 0.001** = 0.754 |
| ED 3c | Sequence duration (young) | One Way *repeated measures* ANOVA | Sequence duration (seconds) | Day (within subjects) | $F_{(2, 22)} = 3.29$ | = 0.056 |
| ED 3c | Sequence duration (aged) | One Way *repeated measures* ANOVA | Sequence duration (seconds) | Day (within subjects) | $F_{(2, 30)} = 7.53$ | **< 0.05** |
| 3c | Outcome devaluation (young vs aged) | Three Way *mixed* ANOVA | Proportion of presses (%)* *Proportion over total was used to normalise choice performance across groups. | (a) Lever (valued/devalued; within-subjects) (b) Test (outcome 1/outcome 2; within-subjects) (c) Group (young/aged; between subjects) | Main Effect (Lever): $F_{(1,14)} = 17.65$ Lever x Group interaction: $F_{(1,14)} = 4.96$ Main Effect (Test): $F_{(1,14)} = 1$ | **= 0.001** **< 0.05** = 0.334 |
| 3f | Outcome devaluation (aged, interspersed) | Two Way *repeated measures* ANOVA | Presses/min | (a) Test (within-subjects) (b) Lever (within-subjects) | Main Effect (Lever): $F_{(1, 7)} = 0.007$ Main Effect (Test): $F_{(1,7)} = 19.355$ | = 0.933 < 0.05 |
| ED 5a | Instrumental acquisition (aged, interspersed) | One Way *repeated measures* ANOVA | Sequence duration (seconds) | Training (within subjects) | $F_{(11, 77)} = 15.359$ | **< 0.001** |

| Fig. | Analysis | Test | Dep. variable | Factor(s) | Statistic | P value |
|---|---|---|---|---|---|---|
| ED 6b | Outcome devaluation (young vs aged) (interval training) | Three Way *mixed* ANOVA | Proportion of presses (%)*<br><br>*Proportion over total was used to normalise choice performance across groups. | (a) Lever (valued/devalued; within-subjects)<br><br>(b) Test (outcome 1/outcome 2; within-subjects)<br><br>(c) Group (young/aged; between subjects) | Main Effect (Lever): $F_{(1,12)} = 0.008$<br><br>Lever x Group interaction: $F_{(1,12)} = 0.984$<br><br>Main Effect (Test): $F_{(1,12)} = 0.735$ | = 0.931<br><br>< 0.341<br><br>= 0.408 |
| 4b | Instrumental acquisition (post-devaluation) | Two Way *mixed* ANOVA | Lever press rate (press/min) | (a) Training day (within-subjects)<br><br>(b) Group (between subjects) | Main Effect (Training): $F_{(2,42)} = 4.96$<br><br>Training x Group interaction: $F_{(2,42)} = 2.23$ | **< 0.05**<br><br>= 0.082 |
| 4d | taD1-SPN density (dorsal striatum, overall effect) | One Way ANOVA | Cell density (cells/mm$^2$) | Group (between subjects) | $F_{(2,45)} = 14.67$ | **< 0.001** |
| 4d | taD1-SPN density (dorsal striatum, simple effects) | Bonferroni *post hoc* | Cell density (cells/mm$^2$) | — | Simple Effects:<br>Young vs. Aged<br>Young vs. Yoked<br>Aged vs. Young<br>Aged vs Yoked | <br>**< 0.001**<br>**< 0.001**<br>**< 0.001**<br>> 0.05 |
| 4d | taD2-SPN density (dorsal striatum, overall effect) | One Way ANOVA | Cell density (cells/mm$^2$) | Group (between subjects) | $F_{(2,45)} = 13.42$ | **< 0.001** |
| 4d | taD2-SPN density (dorsal striatum, simple effects) | Bonferroni *post hoc* | Cell density (cells/mm$^2$) | — | Simple Effects:<br>Young vs. Aged<br>Young vs. Yoked<br>Aged vs. Young<br>Aged vs Yoked | <br>**< 0.001**<br>**= 0.001**<br>**< 0.001**<br>> 0.05 |
| 5b | hM3Dq-pMAPK CNO effect | One Way *repeated measures* ANOVA | Cell density (cells/mm$^2$) | — | Main Effect (Area): $F_{(1,3)} = 124.11$ | **< 0.01** |
| ED 9b | Instrumental acquisition (pre-devaluation) | Two Way *mixed* ANOVA | Lever press rate (press/min) | (a) Training day (within-subjects)<br><br>(b) Group (between subjects) | Main Effect (Training): $F_{(13,234)} = 24.80$<br><br>Mauchly's sph. test: $p < 0.001$<br>Greenhouse Geisser correction: $F_{(2.61,46.95)} = 24.80$<br><br>Training x Group interaction: $F_{(2.61,46.95)} = 1.53$ | <br><br>**< 0.001**<br><br>= 0.221 |
| ED 9c | Sequence length | Two Way *mixed* ANOVA | Presses/sequence | (a) Day (within-subjects)<br><br>(b) Group (between subjects) | Main Effect (Day): $F_{(1, 18)} = 58.37$<br><br>Day x Group interaction: $F_{(1, 18)} = 1.93$ | **< 0.001**<br><br>= 0.181 |
| ED 9d | Sequence duration | Two Way *mixed* ANOVA | Sequence duration (seconds) | (a) Day (within-subjects)<br><br>(b) Group (between subjects) | Main Effect (Day): $F_{(1, 18)} = 0.29$<br><br>Day x Group interaction: $F_{(1, 18)} = 0.374$ | = 0.595<br><br>= 0.548 |
| ED 9e | Concatenation | Two Way *mixed* ANOVA | Concatenated sequences (%) | (a) Day (within-subjects) | Main Effect (Day): $F_{(1, 18)} = 31.41$ | **< 0.001** |

| Fig. | Analysis | Test | Dep. variable | Factor(s) | Statistic | P value |
|---|---|---|---|---|---|---|
| | | | | (b) Group (between subjects) | Day x Group interaction: $F_{(1, 18)} = 1.76$ | = 0.201 |
| **Fig.** | **Analysis** | **Test** | **Dep. variable** | **Factor(s)** | **Statistic** | **P value** |
| 5d | Instrumental acquisition (post-devaluation) | Two Way *mixed* ANOVA | Lever press rate (press/min) | (a) Training day (within-subjects)<br><br>(b) Group (between subjects) | Main Effect (Training): $F_{(3,54)} = 3.103$<br><br>Training x Group interaction: $F_{(3,54)} = 0.565$ | **< 0.05**<br><br>= 0.640 |
| 5f | Outcome devaluation (Sham vs DREADDs) | Two Way *mixed* ANOVA | Lever press rate (press/min | (a) Test (chow/grain; within-subjects)<br><br>(b) Group (between subjects) | Main Effect (Test): $F_{(1,18)} = 7.42$<br><br>Test x Group interaction: $F_{(1,18)} = 4.82$ | **< 0.05**<br><br>**< 0.05** |
| 5j | taD1-SPN density (pDMS, simple effect) | One Way ANOVA | Cell density (cells/mm$^2$) | Group (between subjects) | $F_{(1,38)} = 0.22$ | = 0.883 |
| 5j | taD2-SPN density (pDMS, simple effect) | One Way ANOVA | Cell density (cells/mm$^2$) | Group (between subjects) | $F_{(1,38)} = 7.51$ | **< 0.05** |

**Extended Data Figures**

### Extended Data Fig. 1

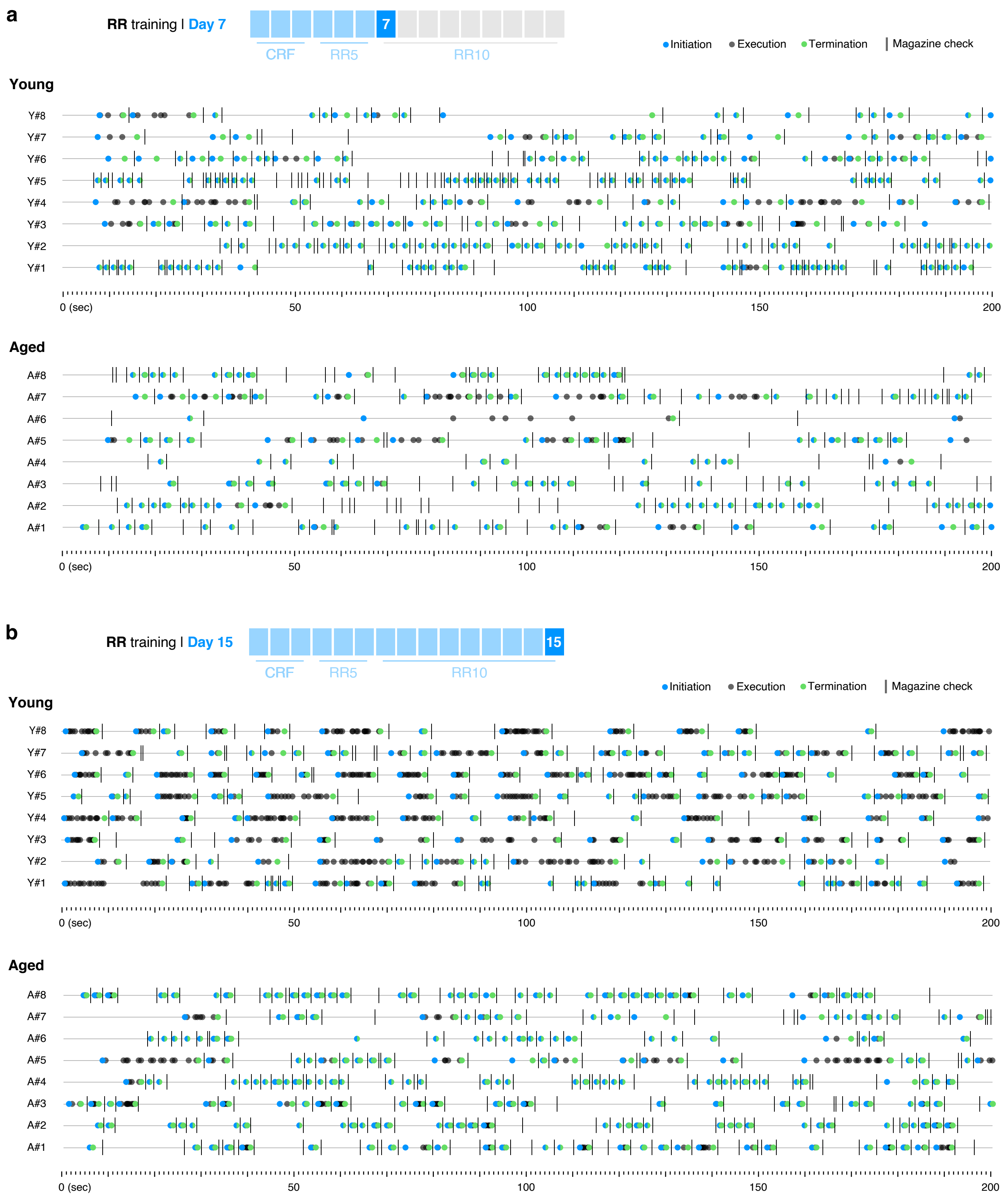


**Extended Data Fig. 1 | Sequential action performance in young and aged mice across single action → outcome training.** Performance was distributed over initiation, execution and termination categories based on lever press and magazine entry interspersing (see Fig. 1a). **a-b**, Event-time diagrams showing linearly organised lever press sequences produced by all young and all aged mice on training days 7 (a) and 15 (b). Top panels show the stage of training data were extracted from. Animal ID numbers are shown on the left.

Extended Data Fig. 2

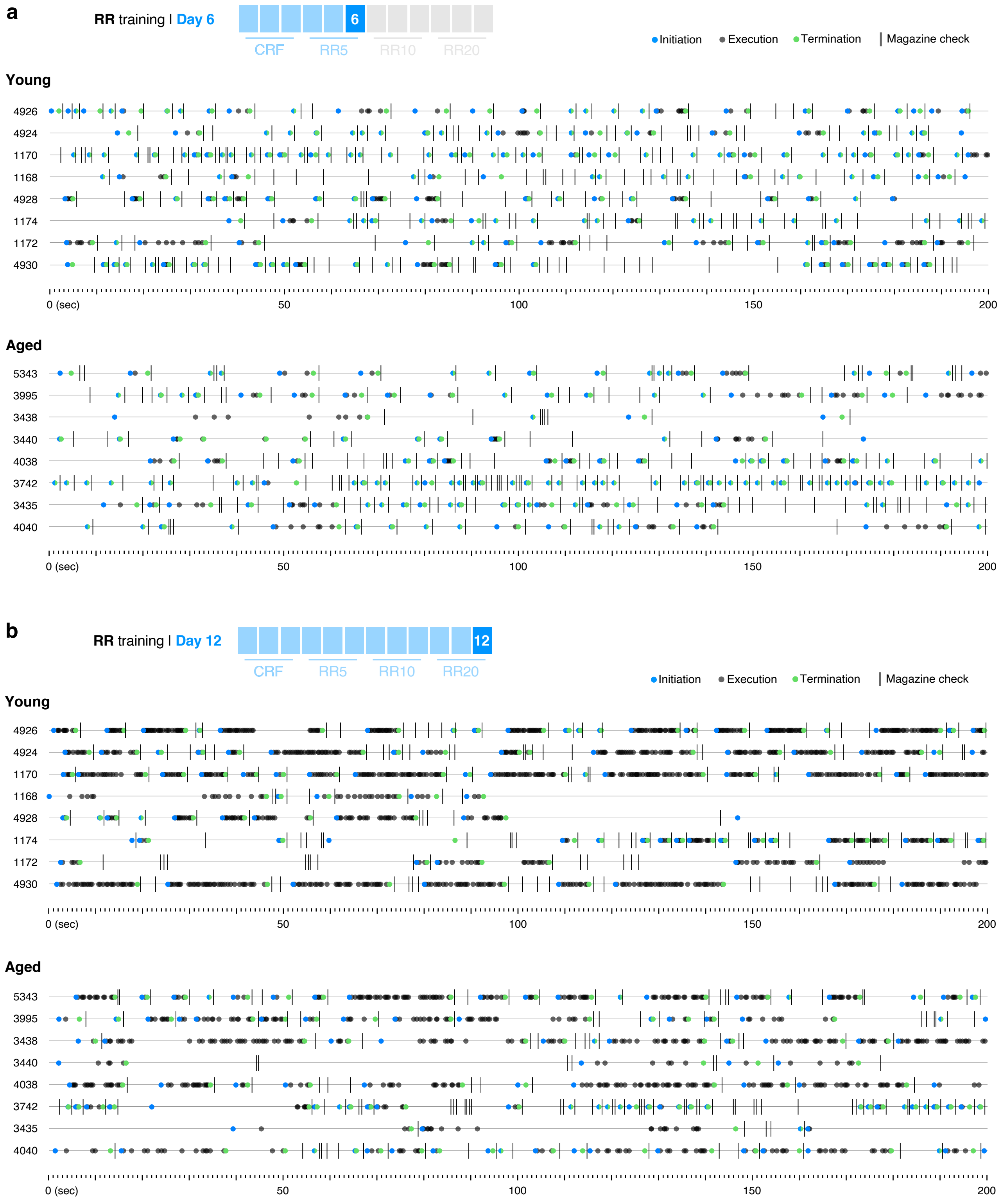


**Extended Data Fig. 2 | Sequential action performance in young and aged mice across dual action → outcome training (ratio schedules).** Performance was distributed over initiation, execution and termination categories based on lever press and magazine entry interspersing (see Fig. 1a). **a-b**, Event-time diagrams showing linearly organised lever press sequences produced on one lever by all young and all aged mice on training days 6 (a) and 12 (b). Top panels show the stage of training data are extracted from. Animal ID numbers are shown on the left.

### Extended Data Fig. 3

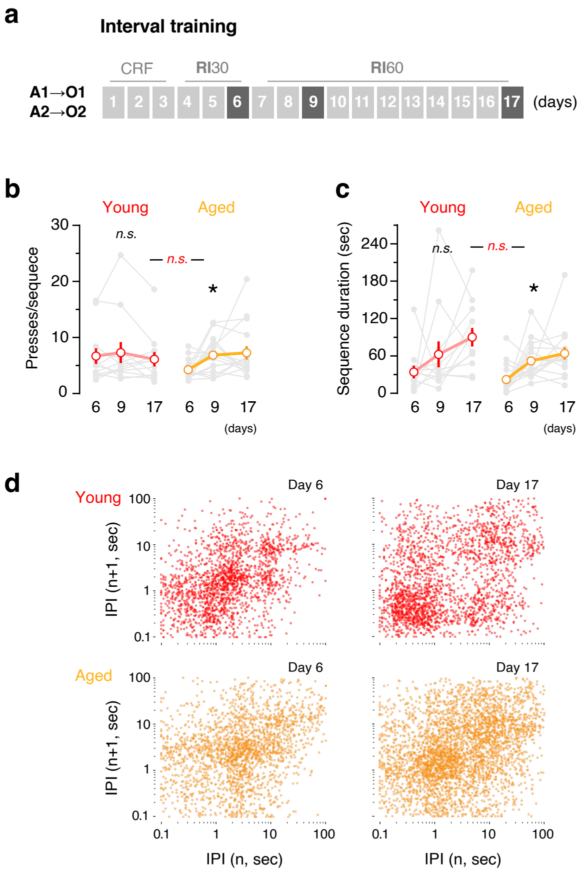


**Extended Data Fig. 3 | Sequential action is altered in both young and aged mice when multiple A→O contingencies are trained under interval schedules of reinforcement. a**, Mice were trained on two different action → outcome contingencies (A1→O1; A2→O2) under continuous reinforcement (CRF) followed by increasing random interval (RI) reinforcement schedules for 17 days. Contingencies were trained in separate sessions each day. **b-c**, Average (mean ± SEM) presses per sequence (b) and sequence duration (c) by each young and aged mouse on days 6, 9 and 17. Grey traces are individual mice (6-8 mice, 2 sessions per mouse). **d**, Return maps showing the temporal relationships between consecutive lever presses at different stages of A→O training. Each dot corresponds to a pair of consecutive IPIs and includes the time delay of each press to its preceding (n [x]) and succeeding (n + 1 [y]) press in the sequence (in seconds [log10]). *, overall/simple effect (black) and interaction (red). n.s., not significant (Supplementary Table 1).

Extended Data Fig. 4

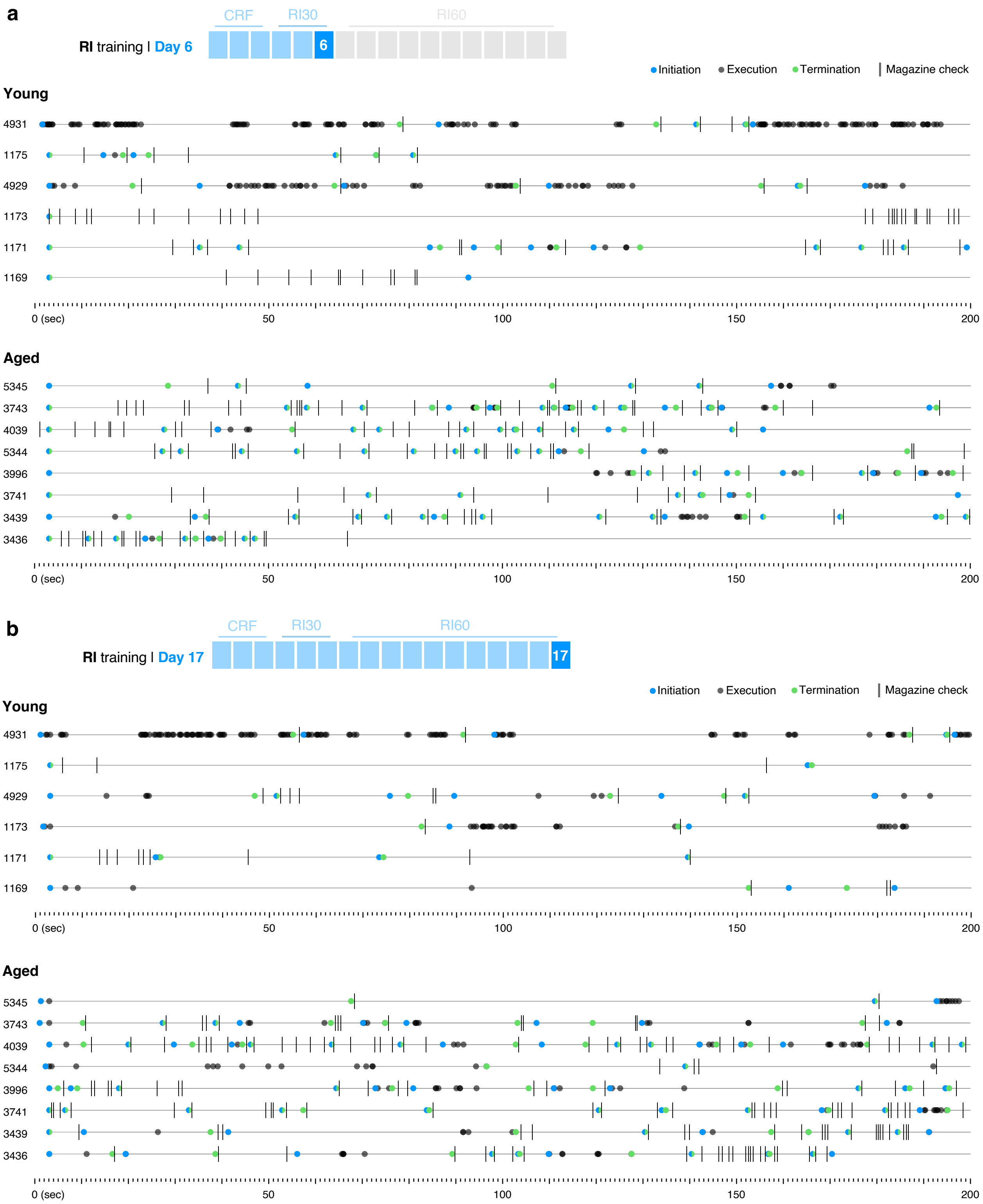


**Extended Data Fig. 4 | Sequential action performance in young and aged mice across dual action → outcome training (interval schedules).** Performance was distributed over initiation, execution and termination categories based on lever press and magazine entry interspersing (see Fig. 1a). **a-b**, Event-time diagrams showing linearly organised lever press sequences produced on one lever by all young and all aged mice on training days 6 (a) and 17 (b). Top panels show the stage of training data are extracted from. Animal ID numbers are shown on the left.

Extended Data Fig. 5

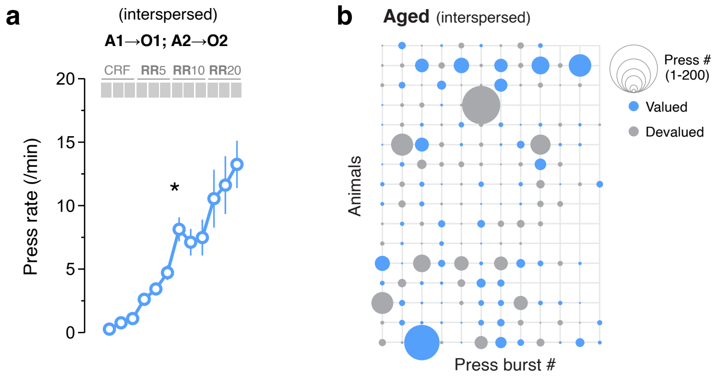


**Extended Data Fig. 5 | Interspersed training of two A→O contingencies does not improve goal-directed vigour in aged mice. a**, Lever presses per minute (press rate) displayed throughout instrumental training (averaged A1 and A2; 8 mice per group). Outcome-specific devaluation test data is shown in Figs. 3c and e. **b**, Bubble plot representing the response vigour exerted on devalued- (grey) and valued- (blue) paired levers throughout the choice test. Each line represents performance by one mouse in one session (two lines per mouse, 8 mice per group). Press bursts are defined as all performance within a 20 second interval, and are organised chronologically. *, overall/simple effect (black). n.s., not significant (Supplementary Table 1).

Extended Data Fig. 6

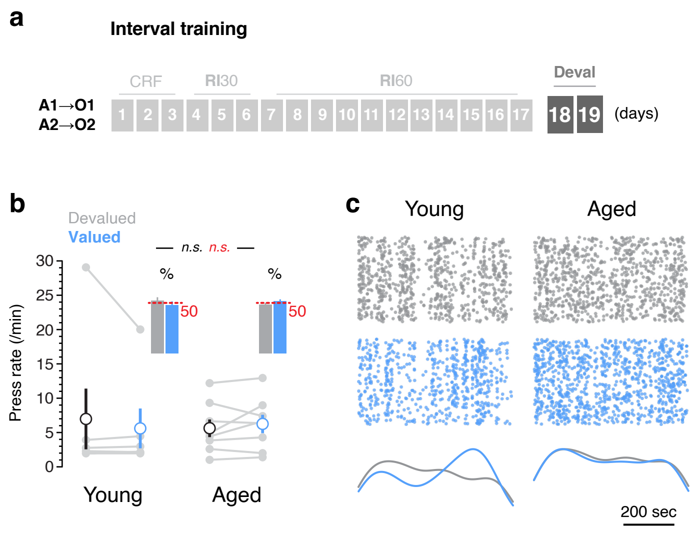


**Extended Data Fig. 6 | Mice fail to develop goal-direction when multiple A→O contingencies are trained under interval schedules of reinforcement, irrespective of age. a**, Top: two separate action → outcome contingencies (A1→O1 and A2→O2) were trained sequentially (in separate sessions) under continuous (CRF) and increasing random interval (RI30 and RI60) reinforcement schedules. Assessment of A→O contingency learning was conducted through outcome-specific devaluation tests (Deval), where mice were sated for 1 hr on one or the other outcome (i.e., O1 or O2) prior to a choice test (A1 versus A2, see Fig. 3A). **b**, Mean ± SEM of press rates on the lever coding for the sated (devalued, grey) and nonsated (valued, blue) outcomes during the choice tests after A →O training. Pairs of data points (light grey) are individual trials (n = 6-8 mice per group; each mouse was exposed to one trial per day on two consecutive days; see Online Methods for details). Insets show the percentage of presses devoted to valued-paired or devalued-paired levers over the total presses in the tests (dashed red line denotes 50%). **c**, Scatter plot of all lever presses performed by the full cohort of young and aged mice during the 10-min choice tests after A→O training. Performance is distributed over devalued- (grey) and valued- (blue) paired levers. Corresponding Kernel density curves of each lever type are plotted at the bottom.

#### Extended Data Fig. 7

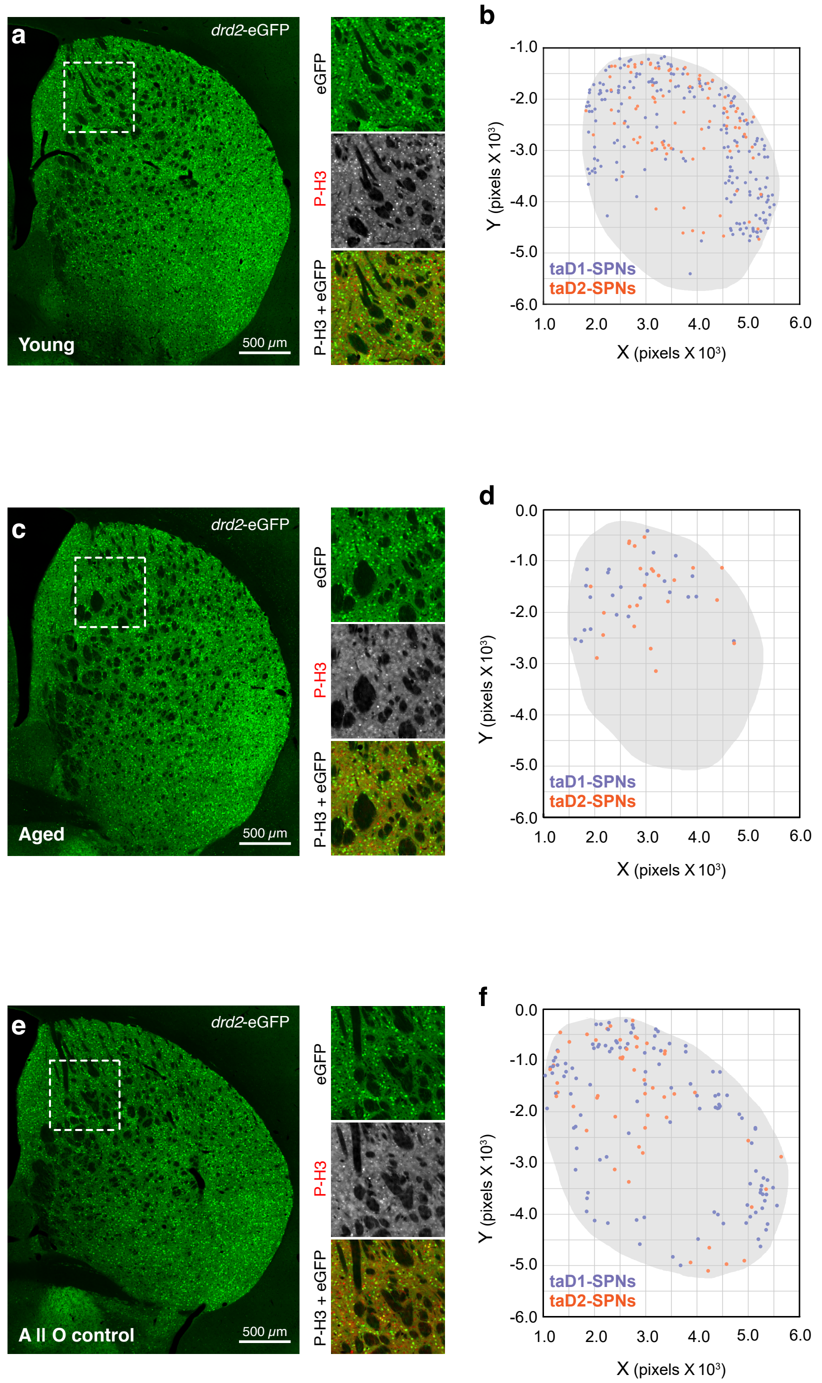


**Extended Data Fig. 7 | Mapping of transcriptionally active SPNs of each type in the striatum. a**, **c**, **e**, Mapping was performed on high resolution (>6,000 px2) high bit depth (16 bit) confocal mosaics of superimposed *drd2*-eGFP (Ch01) and phospho-histone H3 (P-H3; Ch02) signals as previously25. **b**, **d**, **f**, Digitised maps of each image were obtained by plotting the XY coordinate of the centroid of each classified particle (in pixels) in a bidimensional space. Data shown is one example map of a young (a-b), aged (c-d) and control (e-f) subject.

### Extended Data Fig. 8

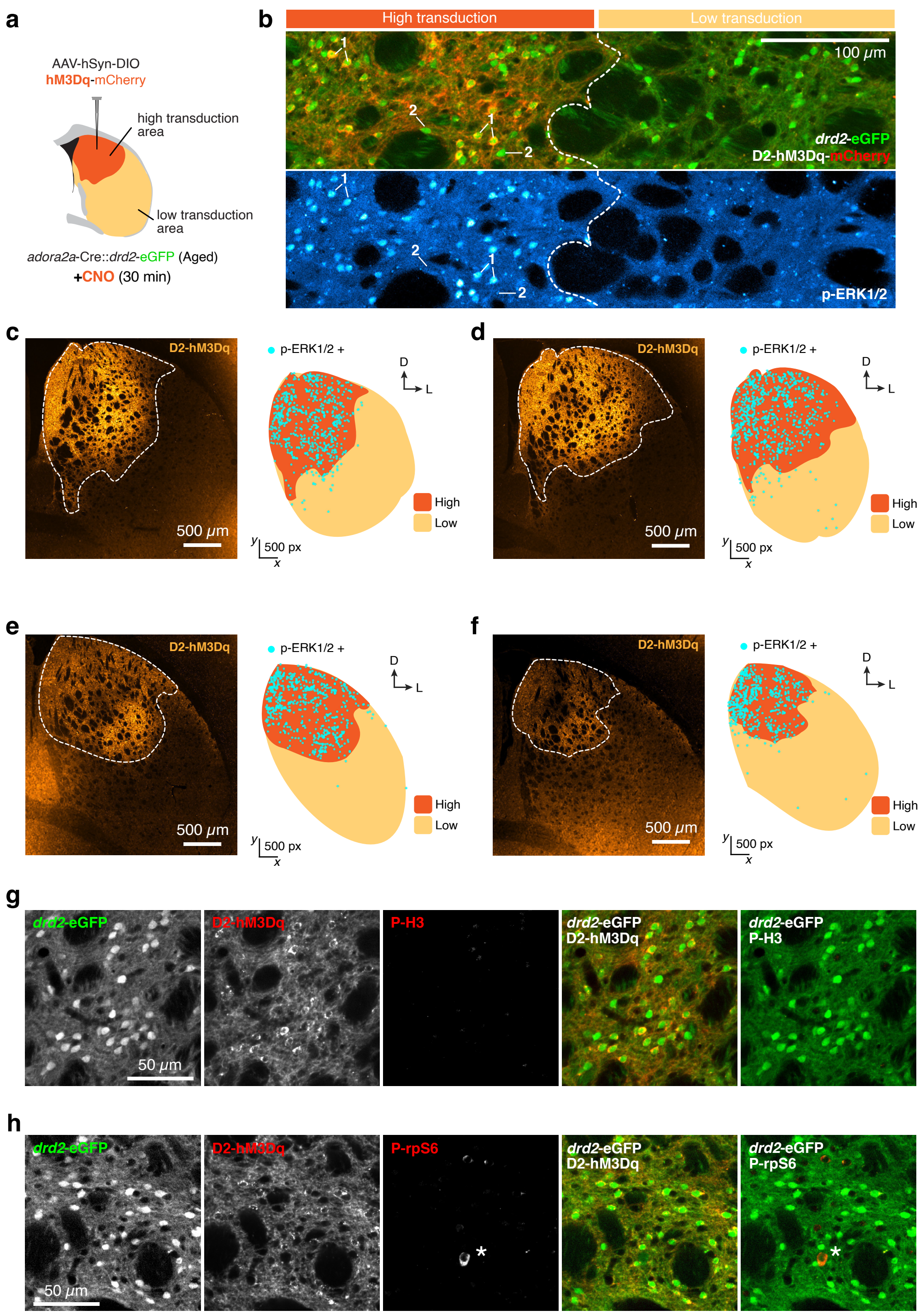


**Extended Data Fig. 8 | Chemogenetic stimulation of intracellular signalling in aged D2-SPNs of the striatum. a**, An AAV carrying Cre-inducible hM3Dq-mCherry was bilaterally microinjected in the posterior dorsomedial striatum (pDMS) of two aged adora2a-Cre x drd2-eGFP hybrid mice (D2-hM3Dq). Three weeks after surgery, mice were given a single injection of CNO (3 mg/kg, i.p.). **b**, Wide-field confocal micrographs showing collocalisation of drd2-eGFP and D2-hM3Dq (top) and corresponding p-ERK1/2 distribution (bottom) in continuous striatal areas with and without viral transduction (high vs. low transduction). **c-f**, Activated neurons in response to CNO were mapped in the pDMS of each hemisection (c-f) through immunodetection of ERK1/2 phosphorylation (Thr202/Tyr204 residues; p-ER1/2) 30 minutes after the injection. Note how activated neurons clustered only in virally transduced areas (distribution in high vs los transduction spaces). Panel b shows how p-ERK1/2 expression was restricted to drd2-GFP(+) (green), hM3Dq(+) (red) neurons within high transduction zones (type 1 cell). Drd2-GFP neurons that did not integrate the viral construct (drd2-GFP(+), hM3Dq(-)) did not express p-ERK1/2, even if located within high transduction zones (type 2 cell). **g-h**, Confocal micrographs showing the absence of phospho-histone H3 (g) and phospho-ribosomal protein S6 (h) upon CNO injection, suggesting that cAMP-dependent signalling remained silent in D2-hM3Dq-expressing neurons[37,38]. Asterisk indicates a putative cholinergic interneurons[50].

Extended Data Fig. 9

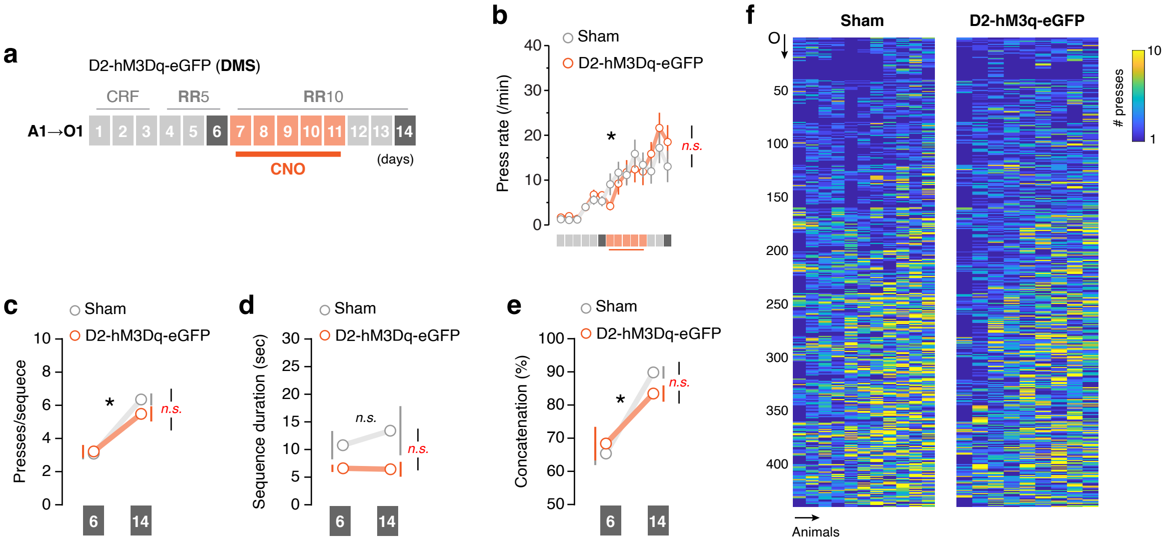


**Extended Data Fig. 9 | CNO treatment in D2-hM3Dq-eGFP hybrid mice did not affect performance during training, which showed some sequential structure. a**, Mice were trained on a single action (A1) → outcome (O1) contingency under continuous reinforcement (CRF) followed by increasing random ratio (RR) reinforcement schedules for 14 days. CNO (3 mg/kg, i.p.) was injected on days 7-11 30 mins prior to training. **b**, Lever presses per minute (press rate, mean ± SEM) displayed throughout instrumental training. **c-d**, Average (mean ± SEM) presses per sequence (c) and sequence duration (d) on days 6 and 14. **e**, Average percentage (mean ± SEM) of lever press concatenation defined as number of sequences with ≥ 2 presses over total. **f**, Heatmap showing the concatenation extent of lever press sequences preceding each outcome, organised chronologically from start to end of training. Nine-to-eleven mice per group; *, overall/simple effect (black) and interaction (red). n.s., not significant (Supplementary Table 1).
